# Regulation of tmTNF-α Processing by FRMD8 in Triple-Negative Breast Cancer Metastasis: Insights into Molecular Pathway Dynamics

**DOI:** 10.1101/2024.09.26.615107

**Authors:** Jun Xu, Xiaoyu Yang, Peng Shu, Wei Wang, Haibo Wu, Zhe Wang

**Author notes:** Address correspondence to: Haibo Wu, MD., Department of Pathology, the First Affiliated Hospital of USTC, Division of Life Sciences and Medicine, University of Science and Technology of China, lujiang road street, 230001 Hefei, Anhui, China. Zhe Wang, MD. State Key Laboratory of Cancer Biology, Department of Pathology, Xijing Hospital and School of Basic Medicine, Fourth Military Medical University, 169 Changle West Road, Xi’an 710032, China. These authors contributed equally: Jun Xu, Xiaoyu Yang.

## Abstract

**Purpose:** Breast cancer remains the leading cause of cancer-related mortality among women worldwide, with late-stage diagnoses prevalent in China resulting in significantly lower survival rates. This study focuses on identifying genes implicated in breast cancer metastasis, highlighting the role of Tumor Necrosis Factor-alpha (TNF-α) and its forms—transmembrane (tmTNF-α) and soluble (sTNF-α).

**Experimental Design:** TNF-α is crucial for activating NF-κB pathways that regulate genes involved in cell adhesion, migration, and immune evasion, all essential for cancer metastasis. We conducted comprehensive analyses of FRMD8, a member of the FERM domain-containing proteins, as a significant regulator of tmTNF-α. Through integrative multi-omics and cellular functional studies, the relationship between FRMD8, iRhom2, and ADAM17 was assessed in the context of breast cancer metastasis.

**Results:** Our findings reveal that FRMD8 forms a complex with iRhom2 and ADAM17, enhancing the stability and sheddase activity of ADAM17, which is vital for the release of TNF-α. The absence of FRMD8 leads to decreased ADAM17 activity, increasing the availability of tmTNF-α and potentially promoting metastasis. This effect suggests that FRMD8 is a key modulator of TNF-α processing.

**Conclusions:** This study explores how FRMD8 influences TNF-α processing and the metastatic behavior of breast cancer, providing insights into molecular dynamics that could guide future therapeutic strategies to improve outcomes in breast cancer patients.

## Introduction

Breast cancer remains the most prevalent cancer among women globally, with over one million new cases diagnosed annually and at least 400,000 fatalities each year, accounting for 14% of all cancer-related deaths [1]. In China, five-year survival rates for patients diagnosed at stages I and II have improved to 94% and 83.2%, respectively [2]. However, this rate dramatically decreases to 65% for stage III and plummets to a mere 21% for stage IV patients, with the majority being diagnosed at these later stages [3]. Consequently, identifying genes associated with breast cancer metastasis is critical for advancing treatment strategies and improving patient outcomes.

Tumor Necrosis Factor-alpha (TNF-α), a member of the TNF superfamily, primarily exists in two forms: transmembrane TNF-α (tmTNF-α) and soluble TNF-α (sTNF-α) [4]. TmTNF-α, a transmembrane protein, can initiate the classical NF-κB pathway via TNFR1 and the non-classical pathway via TNFR2 [5]. Through these pathways, tmTNF-α influences the expression of genes related to cell adhesion and migration, such as Intercellular Adhesion Molecule 1 (ICAM-1) and Matrix Metalloproteinases (MMPs), while also modulating immune evasion within the tumor microenvironment and impacting extracellular matrix remodeling [6]. In contrast, sTNF-α, resulting from the cleavage of tmTNF-α, generally binds solely to TNFR1, activating the classical NF-κB pathway. While extensive studies have examined sTNF-α’s relationship with cancer, tmTNF-α remains less explored [7,8]. In breast cancer, it is crucial to determine which genes can modulate TNF-α cleavage and potentially enhance metastatic processes.

FRMD8, a protein belonging to the FERM domain family, has shown a pivotal role in regulating tmTNF-α activity [9]. By anchoring to the intracellular N-terminus of iRhom2, FRMD8 forms a ternary complex with ADAM17, enhancing the latter’s stability and sheddase activity, thereby facilitating the efficient release of inflammatory mediators such as TNF-α [10]. The absence of FRMD8 leads to the degradation of both iRhom2 and mature ADAM17 via the lysosomal pathway, significantly reducing ADAM17’s activity [11]. Also known as TACE (TNF-α Converting Enzyme), ADAM17 enzymatically releases mature cytokines into the extracellular space by cleaving their membrane-bound precursors [12]. However, the specific mechanisms through which FRMD8 may foster breast cancer metastasis via TNF-α release still warrant further investigation.

## Materials and Methods

### Patient Inclusion and Exclusion Criteria

Tumor samples from seven metastatic breast cancer patients and seven non-metastatic breast cancer patients were collected from Ningbo Beilun People’s Hospital. All patients provided informed consent, and the study procedures were approved by the Ethics Committee of the First Affiliated Hospital of USTC, and Ningbo Beilun People’s Hospital, approval number is 2024-103(YS) and 2024-ky372 respectively.

### Cell Lines

The cell lines used in this study include 759, MCF-7, BT-549, MDA-MB-231, and MDA-MB-436. The 759 cell line was extracted from the primary tumor of Brca1^Co/Co; p53^+/Co; MMTV-Cre mice, which has minimal invasive metastatic ability. The MCF-7, BT-549, MDA-MB-231, and MDA-MB-436 cell lines were preserved in our laboratory. All cell lines were cultured in DMEM with 10% FBS at 37°C in a 5% CO₂ atmosphere.

### RNA-seq Sample Preparation

The RNA-seq samples in this study consist of two parts:TNBC samples with metastasis and five TNBC samples without metastasis, provided by Ningbo Beilun People’s Hospital.Breast cancer samples with low FRMD8 expression and three breast cancer samples with high FRMD8 expression, also provided by Ningbo Beilun People’s Hospital.Analysis of Public Datasets to Identify Low-Expression Metastasis-Related Genes. The preliminary screening of metastasis-related genes was conducted using a combination of our hospital’s dataset sequencing and public dataset analysis.

### Hospital Dataset Analysis Procedure

FFPE samples with metastasis and five without metastasis were evaluated by senior pathologists, and tumor cells were scraped for RNA-seq to detect mRNA expression levels in these ten samples.

After obtaining the FASTQ files from RNA-seq, the analysis process followed the tutorial at https://github.com/griffithlab/rnaseq_tutorial. The FASTQ files were first analyzed for sequencing quality using FASTQC software, and adapters were removed using Flexbar software.

The FASTQ files were then aligned to the reference genome hg38 using HISAT2. The SAM files generated by HISAT2 were converted to BAM files using the samtools sort command.

Gene coverage in the BAM files was observed using IGV software, and the alignment efficiency was checked using samtools flagstat and FASTQC. The StringTie software was used to generate transcripts.gtf and gene_abundances.tsv files, and the htseq-count command was used to generate the gene.tsv file (gene expression file).

After obtaining the gene expression files, edgeR was used to normalize the gene expression data and compare the differentially expressed genes between five TNBC patients with metastasis and five without metastasis. Genes with an absolute logFC value greater than 6 and a p-value less than 0.05 were considered significant, resulting in 895 downregulated genes. The sequencing process for three breast cancer samples with low FRMD8 expression and three with high FRMD8 expression followed the same procedure as the FFPE samples.

### Public Dataset Analysis Procedure

The public dataset selected for metastasis in this study was GSE5327, which was downloaded using the GEOquery package. Both the GSE5327 dataset and the corresponding platform information GPL96 were downloaded using the getGEO function.

The expression matrix was obtained using the exprs function on the GSE data, and gene names were annotated using the GPL96 platform. Duplicate gene names were removed from the GSE data, and the maximum expression value for each gene was retained.

The GSE expression data was normalized using the normalize.quantiles function from the preprocessCore package. Patient survival time and outcomes were extracted using the pData function. The average expression of all genes in the GSE data was calculated, and samples with expression higher than the average were set as the high-expression group, while those below the average were set as the low-expression group.

The survfit function was used to construct a prognosis model, and the surv_pvalue function was used to calculate the p-value of the model, obtaining genes with p-values less than 0.05. In this study, 718 survival-related genes were identified.

After obtaining the genes from our hospital dataset analysis and the public dataset analysis, the two gene lists were intersected, resulting in 39 genes that were lowly expressed in primary tumors with metastasis and associated with lung metastasis survival. These 39 genes were designated as the library genes.

### sgRNA Library Screening Process

After identifying 41 library genes, a knockout library was designed for these genes, and library screening was conducted. The library screening process is as follows: The 41 library genes were cross-referenced with mouse genes, resulting in 39 genes in mice. Six sgRNAs were designed for each gene, with the sgRNA sequences provided in the appendix.

The sgRNA sequences were synthesized by Shanghai Kegi Biotechnology. After synthesis, the sgRNAs were packaged into lentiviruses using second-generation lentivirus packaging technology. Briefly, 12 µg of target plasmid was mixed with 9 µg of pSPAX2 and 3 µg of pMD2.G and transfected into 293T cell lines using Lipofectamine 2000. After 48 hours of transfection, the viral supernatant was collected to obtain the library virus.

The titer of the library virus was measured using a single-clone method, and 759 cell lines were infected with the virus at an MOI of 0.1. After 48 hours of infection, the cells were selected with 2 µg/mL puromycin for seven days to obtain the library cells.

The library cells were injected into the mammary glands of ten nude mice, with 1 million cells injected per mouse. After eight weeks, the mice were sacrificed, and the primary tumors, lungs, spleens, livers, kidneys, and brains were collected.

DNA was extracted from these organs and amplified using universal sgRNA primers. Additionally, the sgRNAs corresponding to the 39 library genes were used as forward primers (sgRNA-F1) and library-R1 as reverse primers for PCR to determine which sgRNAs were present in the lungs, spleens, livers, kidneys, and brains.

### Mouse Experiments

The mice used in this study were mainly nude mice, all purchased from Zhejiang Ziyuan Company. The mice were female, 6–8 weeks old, and weighed around 20 grams. The injection method was mammary injection, with 1 million cells injected per mouse. The animal experimental procedures were approved by the Ethics Committee of our hospital, approval number is 202407111005000167828.

### The drug treatment experimental procedures are as follows

Paclitaxel was purchased from Selleckchem, USA (Cat# S1150), and Etanercept was purchased from Pfizer Pharmaceuticals Limited (New York, NY, United States).

759 FRMD8 sgRNA1 + iRhom2 shRNA cell lines were injected into the nude mice at a dose of 1 million cells per mouse.Two weeks after cell injection, the mice were divided into four groups: CON group, Paclitaxel group, Etanercept group, and Paclitaxel + Etanercept group. Each group received intraperitoneal injections of 100 µL DMSO, 5 mg/kg Paclitaxel, 2 mg/kg Etanercept, or 5 mg/kg Paclitaxel + 2 mg/kg Etanercept, respectively, with an injection frequency of once every two weeks.

Eight weeks after cell injection, the mice were sacrificed to examine the volume of primary tumors and metastasis in various organs in each group. Construction of Different Genotypic Cell Lines

### The main cell lines used in this study include the following categories

FRMD8 Knockdown Gene Construction: FRMD8 was targeted by sgRNA1, sgRNA2, and sgRNA3, with sequences provided in the list. An sgRNA sequence with no target was selected as the control (CON). These constructs were cloned into the lentiCRISPR v2 vector and packaged into lentiviruses. Subsequently, 759, MCF-7, BT-549, and MDA-MB-231 cells were divided into CON, FRMD8 sgRNA1, FRMD8 sgRNA2, and FRMD8 sgRNA3 groups. After infecting the cells with the respective lentiviruses for 48 hours, monoclonal selection techniques were used to obtain knockdown cell lines.

Construction of FRMD8 Wild-Type and (Δ150-200) Genotypes: This part was completed by Fenghui Biotechnology in Hunan. The vector backbone was pCDH-EF1-copGFP-T2A-Puro (Addgene, Cat# 72263), with an empty pCDH-EF1-copGFP-T2A-Puro vector as the control.

BRCA1 Knockdown Gene Construction: The BRCA1 shRNA vector was purchased from Santa Cruz Biotechnology (sc-29219), with a non-targeting shRNA as the control vector. BT-549 cell lines were divided into CON and BRCA1 shRNA groups and transfected with the respective vectors using Lipofectamine 2000. Cells were collected 48 hours after transfection for subsequent experiments.

iRhom2 Knockdown and Overexpression Vectors: The iRhom2 shRNA vector was constructed by Shanghai Sangon Biotech, and the iRhom2 overexpression vector was completed by Fenghui Biotechnology in Hunan. The vector backbone was also pCDH-EF1-copGFP-T2A-Puro (Addgene, Cat# 72263).

### Immunohistochemistry (IHC) and Hematoxylin and Eosin (H&E) Staining

The IHC and H&E experiments were performed by Xinle Biotechnology in Hefei. The general steps for IHC are as follows: Mouse or human tissues were fixed in 4% paraformaldehyde overnight and washed with PBS buffer. The tissues were then dehydrated using a graded ethanol series (70%, 80%, 90%, 95%, and 100%).

The tissues were cleared with xylene and embedded in paraffin. Once the paraffin solidified, the tissue blocks were cut into 3–5 μm thick sections and mounted on slides.

The paraffin sections were deparaffinized with xylene, rehydrated through a graded ethanol series, and rinsed with distilled water. Antigen retrieval was performed by high-temperature and high-pressure treatment.

The tissue sections were blocked with BSA to reduce non-specific antibody binding. After incubation with specific primary antibodies, the sections were washed with TBST buffer, followed by the addition of secondary antibodies corresponding to the primary antibody’s host species.DAB staining was used to observe the protein expression intensity of the target genes.

### The general steps for H&E staining are as follows

Tissue fixation and sectioning steps were the same as those for IHC. After obtaining slides with tissue sections, the slides were immersed in hematoxylin stain and rinsed under running water.

The slides were immersed in acidic ethanol solution, rinsed with running water, then immersed in bluing solution, and rinsed again with running water. The slides were immersed in eosin stain, rinsed under running water, and passed through an increasing concentration gradient of ethanol solutions. The tissue sections were cleared with xylene to remove ethanol, and finally, coverslips were applied to the slides for observation under a microscope. After completing the IHC and H&E staining, the slides were scanned using the Pannoramic MIDI machine from 3DHISTECH and stored using CaseViewer scanning software.

### Immunofluorescence

Immunofluorescence staining includes both tissue and cell immunofluorescence staining. The preliminary staining steps were performed by Xinyue Biotechnology in Hefei. The tissue immunofluorescence pre-treatment and primary antibody labeling steps followed the IHC protocol. After primary antibody labeling, tissue sections were labeled with fluorescently tagged secondary antibodies and scanned using the same scanning software as in the IHC experiments.

### The steps for cell immunofluorescence staining are as follows

Place cell coverslips in a 6-well plate and add 1 million cells per well. After the cells adhere, fix them with 4% paraformaldehyde for 15 minutes. Wash the cells with PBS, permeabilize them with 0.1% Triton X-100, and wash again with PBS.

Block the cells with BSA blocking solution and label them with primary antibodies and corresponding secondary antibodies. Finally, add DAPI to stain the nuclei. Mount the cell coverslips onto slides and observe the fluorescence intensity using a Zeiss LSM780 microscope.

### Quantitative PCR (qPCR)

mRNA expression levels were detected using qPCR. The steps are as follows: Place tumor tissue in liquid nitrogen and add an appropriate volume of TRIzol solution. Grind the tissue and collect 500 µL of the supernatant. Add 200 µL of chloroform, shake vigorously, centrifuge, and collect 100 µL of the upper aqueous phase.

Add 100 µL of isopropanol, mix by pipetting, centrifuge, and wash the precipitate twice with 70% ethanol to obtain total mRNA.

Reverse transcribe total mRNA into cDNA using a reverse transcription kit (Takara Bio) and detect the relative expression of genes using a SYBR Green dye kit (Takara Bio). The primers used in this study are listed in the appendix. After obtaining the relative expression levels of the target gene and GAPDH, use GAPDH as the internal control and quantify the relative expression using the 2^–ΔΔCT method.

### Western Blot

Western blotting was performed as follows:Cells were lysed on ice for 30 minutes with RIPA lysis buffer, and an appropriate volume of protein loading buffer was added. The mixture was boiled for 10–15 minutes to prepare the samples.

Samples (30 µg each) were loaded onto pre-cast SDS-PAGE gels. Electrophoresis was performed at 50 V until the dye front reached the separating gel, then at 120 V until the dye front reached the bottom of the gel. Proteins were transferred to a PVDF membrane at a constant current of 400 mA.

The membrane was incubated with the primary antibodies listed in the appendix at 37°C for 4 hours, followed by incubation with the corresponding secondary antibodies (1:5000 dilution).Protein bands were visualized using ECL solution and quantified using ImageJ software. The relative expression level of the target protein was calculated as the ratio of the target protein to GAPDH. Each experiment was repeated three times, and the average value was taken.

### CyTOF and Flow Cytometry Experiments

The CyTOF experiments were performed by Zhejiang ProTing Company. The general steps are as follows:Chop the tissue into small pieces (1–2 mm³) and incubate with a suitable enzymatic cocktail to dissociate the extracellular matrix and release single cells.

Pass the digested tissue suspension through a cell strainer (40–70 µm mesh size) to remove undigested tissue fragments and obtain a single-cell suspension.

Centrifuge the single-cell suspension at low speed, wash, and assess cell viability using viability dyes.Perform a final wash with PBS to remove any remaining debris.Label the cells with appropriate antibodies and use a mass cytometry machine from Fluidigm to detect the content of immune cells.

### T Cell Suppression Assay

Myeloid-derived suppressor cells (MDSCs) were isolated from primary tumor tissue four weeks post-implantation of 759-GFP cells expressing FRMD8 sgRNA1/iRhom2 shRNA. Tumor tissues were digested using digestion buffer for one hour at 37°C. Following digestion, cells were treated with RBC lysis buffer to lyse red blood cells. Single-cell populations were labeled with CD11b and Gr1 antibodies and incubated on ice for 45 minutes. MDSC populations were sorted using a FACS Aria II flow cytometer.

T cells, isolated from wild-type spleen and bone marrow, were labeled with CFSE and combined with anti-CD3 and anti-CD28 antibodies. These T cells were then mixed with MDSCs at ratios of 0:1 or 1:1, with a total of 5.0 × 10LJ cells per well in a 24-well plate. T cell proliferation was assessed by FACS analysis after 96 hours of culture.

### Drug Library Screening and Public Dataset Analysis

A drug library containing 146 FDA-approved drugs was synthesized by Genscript and distributed in a 384-well plate. The distribution of the drugs in the wells is provided in the appendix. Both 759 CON and 759 FRMD8 sgRNA1 cells were plated onto the drug library plate. After 48 hours, 10 µL of MTT solution (5 mg/mL) was added to each well. The plate was incubated at 37°C with 5% CO₂ for 4 hours, after which 100 µL of DMSO was added. The absorbance (OD value) of each well was measured using a microplate reader at a wavelength of 570 nm. Cell sensitivity to the drugs was calculated based on the absorbance ratios.

### Promoter Assay

The promoter sequence of TNF-α, spanning from –1499 to +100, was downloaded from the EPD website and cloned into a pGL-basic vector. This pGL-basic vector or the pGL-TNF vector, along with mBRCA1 cDNA, control DNA, and Renilla DNA, was co-transfected into wild-type and BRCA1-mutant mammary epithelial cells in 24-well plates for 72 hours. After incubation, cells were lysed, and luciferase activity was measured using a luciferase assay system kit (Promega). Quantification of signal intensity was performed by calculating the ratio of Firefly to Renilla luciferase activity.

### Cell Co-culture Experiment

After digesting the primary tumor into a single-cell suspension, MDSCs (Gr1⁺) were obtained using flow cytometry. These cells were mixed with CON and FRMD8 sgRNA1 cells at a ratio of 3:1. The mixed cell culture was placed in a Precise Live Cell Imaging incubator for 48 hours, and cell dynamics were monitored using an Olympus IX71 microscope. The recorded effects were analyzed using the OpenCV-Python package.

### Statistical Analysis

Quantitative data were analyzed using t-tests, while qualitative data were analyzed using chi-square tests. A p-value of less than 0.05 was considered statistically significant.

### Data availability Statement

All data accessed from external sources and prior publications have been referenced in the text and corresponding figure legends. Additional data and requests should be directed to and will be fulfilled by the Lead Contact, Dr. Zhe Wang.

## Results

### Identification of FRMD8 as a Metastasis-Related Gene

To identify metastasis-suppressor genes in breast cancer, we selected five primary tumors with metastasis and five without from our hospital’s data and performed RNA-seq to identify genes downregulated in metastatic tumors, resulting in 936 downregulated genes (Table 1). Similarly, in the GSE5327 dataset, we identified 759 downregulated genes related to lung metastasis (Table 1). Cross-referencing these lists yielded 41 overlapping genes (Figure 1A).

**Figure 1.**
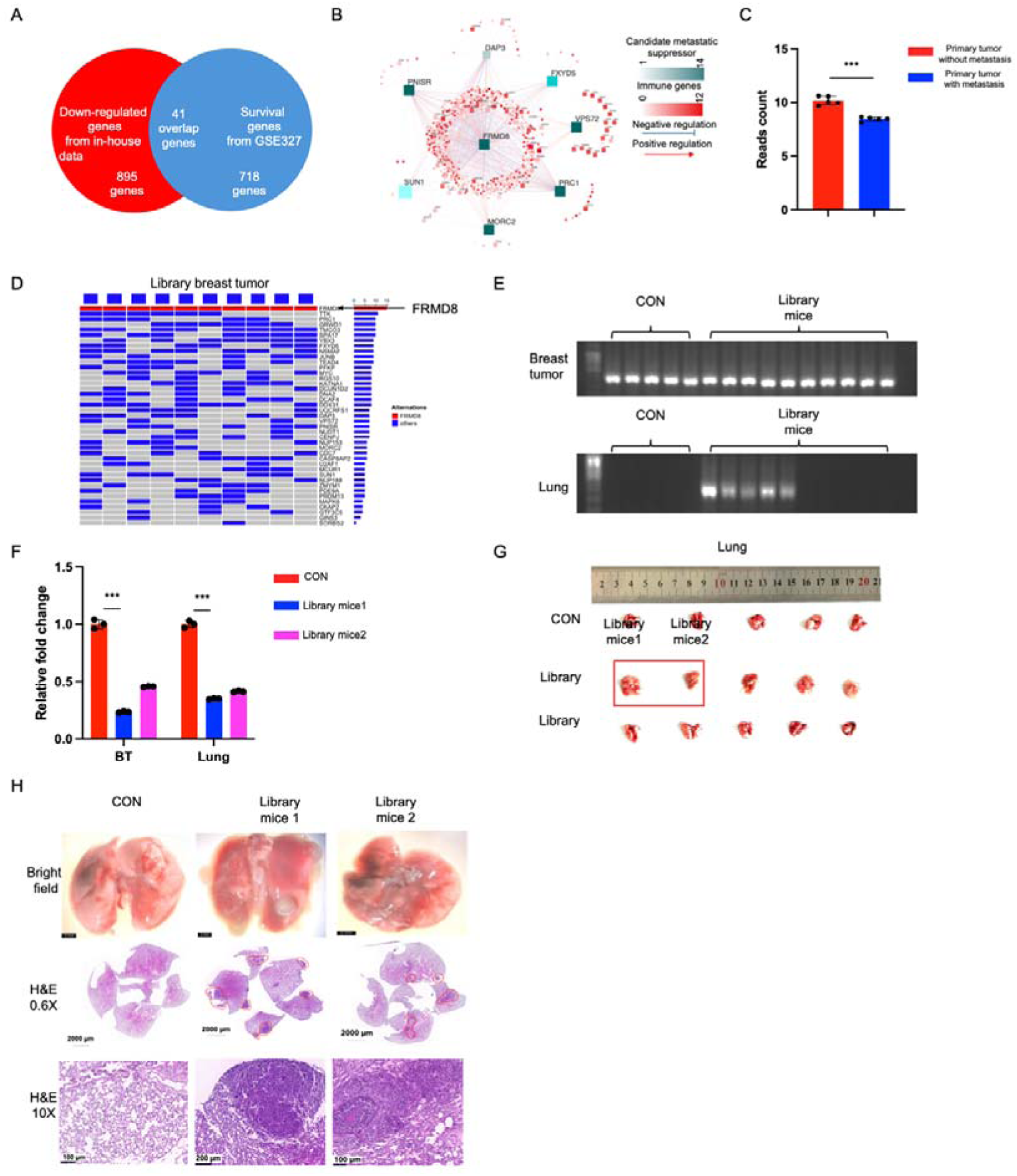
Identification of FRMD8 as a Metastasis-Related Gene Using Library Screening Techniques. (A) Venn diagram showing the overlap of downregulated genes identified in breast cancer patients with metastasis from our hospital’s dataset and the GSE5327 public dataset, resulting in 41 common genes. (B) Transcriptional regulatory network analysis in the GSE5327 dataset illustrating the regulatory relationships among the 41 genes, with FRMD8 highlighted as a key regulator. (C) Reads count analysis showing significantly lower FRMD8 expression in breast cancer tumors with metastasis compared to those without metastasis in our hospital’s dataset. (D) PCR amplification of genomic DNA from control (CON) and library-injected tumors using sgRNA primers, indicating the presence of FRMD8 sgRNA in primary tumors following injection of library cells into the mammary gland. (E) PCR detection of FRMD8 sgRNA presence in primary tumors and lungs, demonstrating that FRMD8 sgRNA is present in the lungs of library mice but not in controls. (F) qPCR analysis of FRMD8 mRNA levels in primary tumors and lungs of control mouse 1, library mouse 1, and library mouse 2, confirming reduced FRMD8 expression in library mice. (G) Gross images of lungs from control mice and library mice, showing visible lung metastasis in library mice. (H) Hematoxylin and eosin (H&E) staining of lung tissues from control mice and library mice, indicating increased lung metastasis in library mice compared to controls.Data are presented as mean ± SD (n = 5 mice per group). Statistical significance was assessed using a two-tailed Student’s t-test. ***P < 0.001.

For these 41 genes, we designed a small sgRNA knockout library (Table 1) and used transcriptional regulatory network analysis in the GSE5327 dataset to identify significantly regulated genes (Figure 1B). Master regulator analysis showed that FRMD8 had the most significant regulatory capability (Table 1), and FRMD8 was also downregulated in our hospital’s dataset (Figure 1C). In the TCGA dataset, FRMD8 was confirmed to be downregulated in the TNBC subtype (Figure S1A), which is highly invasive. This, combined with its low expression in metastatic tumors, suggests FRMD8 as a candidate metastasis-suppressor gene.

We further selected the 759 cell line, derived from tumors in BRCA1-deficient mice, which lacks metastatic capability in vivo. We divided the 759 cell line into CON and library groups, injecting five nude mice with 759 CON cells and ten mice with 759 library cells. After eight weeks, we harvested DNA from primary tumors and used primers for the 39 library genes for amplification. All ten library mice tumors contained FRMD8 sgRNA, indicating that low FRMD8 expression may support breast cancer proliferation (Figure 1D).

PCR analysis of primary tumors and lungs from these mice showed FRMD8 sgRNA in the lungs of five library mice but not other library genes (Figure 1E). Two of these mice had significantly enlarged spleens (Figure S1B), and despite no visible tumors in their brains (Figure S1C), livers (Figure S1D), or kidneys (Figure S1E), PCR detected FRMD8 sgRNA in these organs, suggesting a multi-organ metastatic capability (Figure S1F).

Further H&E staining confirmed increased metastasis in these mice compared to controls (Figures 1F, 1G). Library mouse 1 showed visible lung metastasis (Figure 1G), and H&E staining confirmed increased lung metastasis in library mice compared to CON (Figure 1H). No visible metastasis was found in the liver, spleen, kidneys, or brains of CON or library mice, suggesting that low FRMD8 expression primarily drives lung metastasis.

### Low Expression of FRMD8 Promotes Multi-Organ Metastasis in Mice

We selected seven primary tumors with metastasis and seven without from our hospital’s data and assessed FRMD8 expression at the mRNA and protein levels using qPCR and Western blot. Results showed significantly reduced FRMD8 expression in metastatic tumors (Figures 2A, 2B). The GSE5327 dataset also indicated a negative correlation between FRMD8 and patient survival in lung metastasis (Figure S2A).

**Figure 2.**
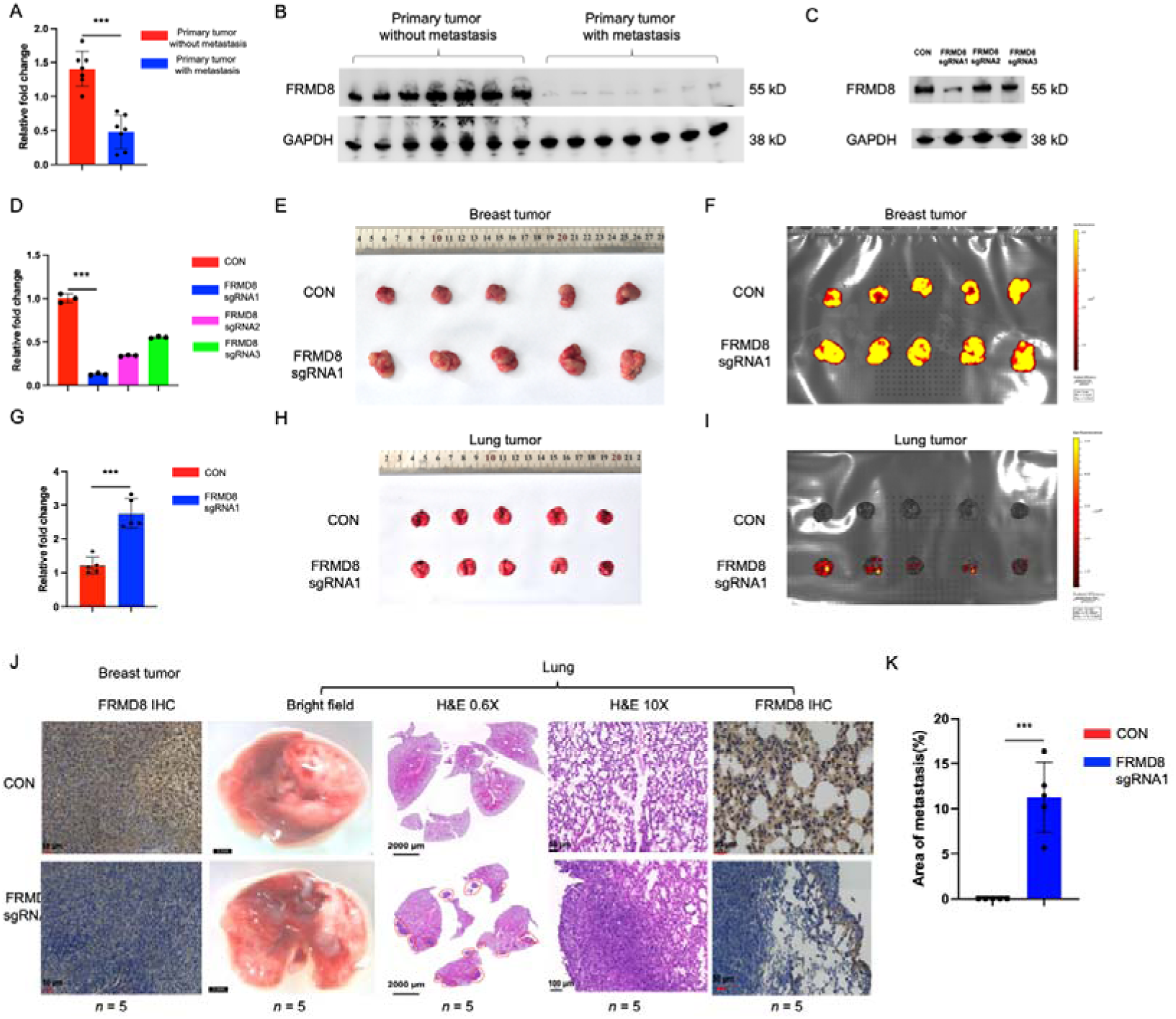
Low Expression of FRMD8 Promotes Multi-Organ Metastasis in Mice. (A) qPCR analysis of FRMD8 mRNA expression levels in primary tumors from seven cases with metastasis and seven cases without metastasis from our hospital’s dataset, showing significantly reduced FRMD8 expression in metastatic tumors. (B) Western blot analysis of FRMD8 protein levels in primary tumors from seven cases with metastasis and seven cases without metastasis, confirming decreased FRMD8 expression in metastatic tumors. (C) Western blot detection of FRMD8 protein levels in 759 cells transfected with control (CON), FRMD8 sgRNA1, FRMD8 sgRNA2, and FRMD8 sgRNA3, demonstrating effective knockdown by sgRNA1. (D) qPCR analysis of FRMD8 mRNA levels in 759 CON, 759 FRMD8 sgRNA1, 759 FRMD8 sgRNA2, and 759 FRMD8 sgRNA3 cells, confirming significant knockdown by sgRNA1. (E) Images showing changes in primary tumor volume in mice injected with 759 CON and 759 FRMD8 sgRNA1 cells, with larger tumors observed in the FRMD8 sgRNA1 group. (F) In vivo imaging of GFP signals in primary tumors from 759 CON and 759 FRMD8 sgRNA1 groups, indicating increased tumor growth in the FRMD8 sgRNA1 group. (G) Statistical analysis of primary tumor volumes in 759 CON and 759 FRMD8 sgRNA1 groups, showing a significant increase in the FRMD8 sgRNA1 group. (H) Gross images of lungs from mice injected with 759 CON and 759 FRMD8 sgRNA1 cells, revealing visible lung metastases in the FRMD8 sgRNA1 group. (I) Quantification of GFP signal intensity in lungs from 759 CON and 759 FRMD8 sgRNA1 groups, indicating increased metastasis in the FRMD8 sgRNA1 group. (J) Immunohistochemical staining of FRMD8 expression in primary tumors and lung metastatic tumors from 759 CON and 759 FRMD8 sgRNA1 groups, confirming reduced FRMD8 expression in the FRMD8 sgRNA1 group. (K) Statistical analysis of lung metastasis incidence in 759 CON and 759 FRMD8 sgRNA1 groups, showing a significant increase in the FRMD8 sgRNA1 group.Data are presented as mean ± SD (n = 5 mice per group). Statistical significance was assessed using a two-tailed Student’s t-test. ***P < 0.001.

To further investigate FRMD8’s role in metastasis, we introduced FRMD8 sgRNA1, sgRNA2, and sgRNA3 into the 759 cell line. FRMD8 sgRNA1 significantly reduced FRMD8 protein levels (Figure 2C, Figure S2B), and qPCR confirmed similar results (Figure 2D). Injecting 759 CON and FRMD8 sgRNA1 cells into the mammary glands of nude mice, we observed increased primary tumor size in the FRMD8 sgRNA1 group (Figures 2E, 2F), with statistically significant differences in tumor volume (Figure 2G).

In addition to enhanced tumor proliferation, we observed visible lung metastasis in the FRMD8 sgRNA1 group (Figure 2H). Although the lung area did not significantly differ between the CON and FRMD8 sgRNA1 groups (Figure S2C), metastasis intensity was significantly higher in the FRMD8 sgRNA1 group (Figure 2I). IHC showed significantly reduced FRMD8 expression in primary and lung metastatic tumors in the FRMD8 sgRNA1 group (Figure 2J), with statistically significant increased lung metastasis (Figure 2K).

For other organs, no visible metastasis was observed in the liver, spleen, brain, or kidneys of FRMD8 knockdown mice, but in vivo imaging confirmed increased metastasis in these organs in the FRMD8 sgRNA1 group (Figures S2D–G). Literature review revealed that FRMD8’s function requires the FERM domain (Figure S2H). We designed a 150–200 amino acid deletion in FRMD8 and introduced it into the 759 cell line, resulting in slightly reduced molecular weight (Figure S2I). Overexpression of FRMD8(Δ150–200) significantly increased in vivo metastasis capability of the 759 cell line (Figures S2J–L).Combining these data, we concluded that low FRMD8 expression significantly promotes multi-organ metastasis in mice.

### Knockdown of FRMD8 Increases tmTNF-**α** Expression and Activates Metastasis-Related Pathways

To further elucidate the mechanism by which FRMD8 influences metastasis, we selected six breast tumor samples from our hospital’s data—three with high FRMD8 expression and three with low expression. qPCR results confirmed the effective reduction of FRMD8 in the low-expression group (Figure S3A). Pathway enrichment analysis revealed significant enrichment in protein localization to the membrane and macromolecular complex pathways in FRMD8-low tumors (Figures 3A, S3B, S3C; Table 2). Given FRMD8’s role in membrane protein cleavage, we hypothesized its involvement in cytokine secretion.

**Figure 3.**
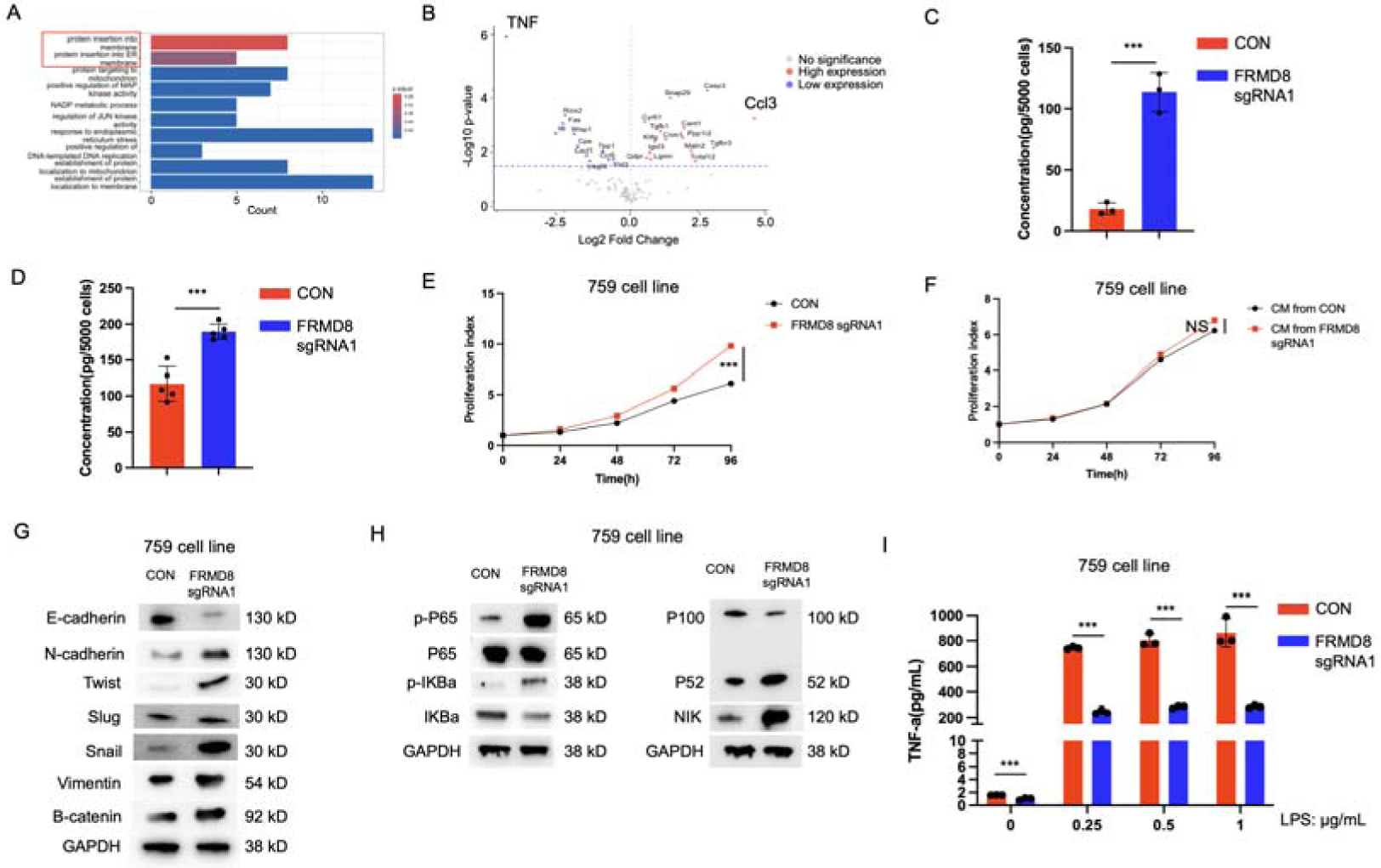
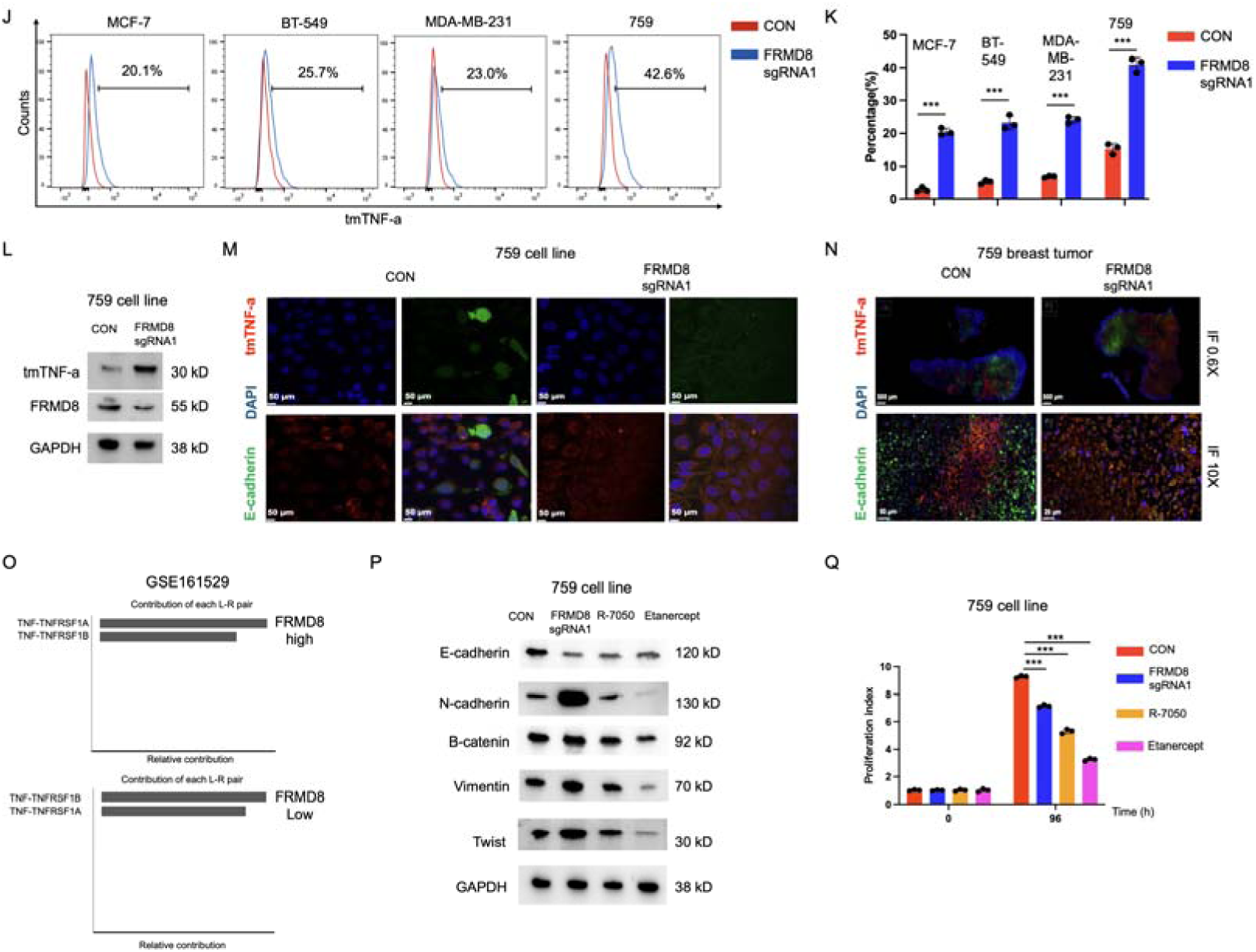
Knockdown of FRMD8 Increases tmTNF-α Expression and Activates Metastasis-Related Pathways. (A) Pathway enrichment analysis of tumors with low and high FRMD8 expression, showing significant enrichment in protein localization to the membrane and macromolecular complex pathways in FRMD8-low tumors. (B) Cytokine array analysis of supernatants from 759 CON and 759 FRMD8 sgRNA1 cells, indicating decreased TNF-α and increased CCL3 levels in FRMD8 knockdown cells. (C) ELISA measurement of CCL3 levels in the supernatant of MDA-MB-436 CON and FRMD8 sgRNA1 cells, confirming increased CCL3 secretion upon FRMD8 knockdown. (D) qPCR analysis of CCL3 expression in primary tumors derived from 759 CON and 759 FRMD8 sgRNA1 cells, showing elevated CCL3 levels in the FRMD8 sgRNA1 group. (E) Cell proliferation assay of 759 CON and 759 FRMD8 sgRNA1 cells, demonstrating increased proliferation in FRMD8 knockdown cells. (F) Cell proliferation assay of 759 cells treated with supernatants from 759 CON and 759 FRMD8 sgRNA1 cells, showing no significant change, indicating that the effect is cell-autonomous. (G) Western blot analysis of EMT markers (E-cadherin and N-cadherin) in 759 CON and 759 FRMD8 sgRNA1 cells, showing EMT activation upon FRMD8 knockdown. (H) Western blot analysis of classical and non-classical NF-κB pathway activation in 759 CON and 759 FRMD8 sgRNA1 cells, indicating increased pathway activation with FRMD8 knockdown. (I) ELISA measurement of TNF-α levels in the supernatant of 759 CON and 759 FRMD8 sgRNA1 cells treated with different concentrations of LPS, showing reduced TNF-α secretion upon FRMD8 knockdown. (J) Flow cytometry analysis of surface tmTNF-α expression in MCF-7, BT-549, MDA-MB-231, and 759 cells transfected with CON and FRMD8 sgRNA1, revealing increased tmTNF-α levels with FRMD8 knockdown. (K) Statistical analysis of flow cytometry data from (J), confirming significant increases in tmTNF-α expression. (L) Western blot analysis of tmTNF-α and FRMD8 expression in 759 CON and 759 FRMD8 sgRNA1 cells, validating increased tmTNF-α upon FRMD8 knockdown. (M) Immunofluorescence staining of tmTNF-α on the cell membrane in 759 CON and 759 FRMD8 sgRNA1 cells, demonstrating elevated tmTNF-α levels in FRMD8 knockdown cells. (N) Immunofluorescence staining of tmTNF-α in primary tumors from 759 CON and 759 FRMD8 sgRNA1 groups, confirming increased tmTNF-α expression in vivo. (O) Analysis of TNF-TNFRSF1B signaling intensity in tissues with high and low FRMD8 expression in BRCA1 mutant primary tumors from the GSE161529 dataset, indicating enhanced TNFR2 signaling with low FRMD8 expression. (P) Western blot analysis of EMT markers in 759 cells treated with the TNFR1 inhibitor R-7050 and the TNFR1/TNFR2 inhibitor Etanercept, showing reduced EMT activation with TNFR2 inhibition. (Q) Cell proliferation assay of 759 cells treated with R-7050 and Etanercept, demonstrating decreased proliferation upon TNFR2 inhibition. Data are presented as mean ± SD. Statistical significance was assessed using a two-tailed Student’s t-test. NS, not significant; ***P < 0.001.

Cytokine array analysis showed decreased TNF-α levels in the supernatant of 759 FRMD8 sgRNA1 cells compared to controls (Figure 3B). However, FRMD8 expression positively correlated with TNF-α levels in both the TCGA-BRCA dataset and the TNBC subtype dataset (Figures S3D, S3E), suggesting that FRMD8 may promote TNF-α transcription but inhibit its secretion.

Moreover, CCL3 levels were significantly increased in the supernatant of FRMD8 sgRNA1 cells (Figures 3B, S3F; Table 2), which was confirmed by ELISA (Figure 3C). CCL3 expression was also elevated in primary tumors derived from 759 FRMD8 sgRNA1 cells (Figure 3D). Since CCL3 is known to recruit myeloid-derived suppressor cells (MDSCs), we propose that FRMD8 knockdown may modulate immune cell functions within the tumor microenvironment.

Cell proliferation assays showed increased growth rates in FRMD8 knockdown cells (Figure 3E), but treatment with the supernatant from FRMD8 sgRNA1 cells did not enhance proliferation in 759 cells (Figure 3F). Similar results were obtained in MCF-7, BT-549, and MDA-MB-231 cell lines, where FRMD8 sgRNA1 showed the most significant knockdown efficiency (Figures 3G–I). In these cell lines, FRMD8 knockdown also led to increased proliferation indices (Figure S3J), suggesting that the effects of FRMD8 are cell-autonomous and rely on cell–cell contact rather than secreted factors.

Regarding epithelial–mesenchymal transition (EMT) markers, N-cadherin expression increased while E-cadherin decreased in 759 FRMD8 sgRNA1 cells (Figure 3G), but these changes were not observed when cells were treated with the supernatant from FRMD8 sgRNA1 cells (Figure S3K). Activation of both the classical and non-classical NF-κB pathways was evident in FRMD8 sgRNA1 cells (Figure 3H) but not induced by the supernatant treatment (Figure S3L). These findings suggest that FRMD8 knockdown affects proliferation, metastasis, and TNF-related pathways independently of soluble factors.

Given FRMD8’s role in cytokine secretion and tmTNF-α cleavage, we assessed TNF-α secretion under different lipopolysaccharide (LPS) concentrations. FRMD8 knockdown decreased TNF-α secretion in the 759 cell line (Figure 3I), and LPS-stimulated TNF-α secretion was reduced in FRMD8 sgRNA1 cells (Figures S3M–O), indicating that FRMD8 knockdown inhibits TNF-α release.

Flow cytometry analysis revealed increased tmTNF-α on the cell membrane in MCF-7, BT-549, MDA-MB-231, and 759 cell lines with FRMD8 knockdown (Figures 3J, 3K), which was confirmed by Western blot (Figure 3L). Immunofluorescence showed elevated tmTNF-α levels both in vitro (Figure 3M) and in primary tumors (Figure 3N), while supernatant treatment did not induce tmTNF-α expression (Figure S3P). Overexpression of FRMD8(Δ150–200) also increased tmTNF-α levels (Figures S3Q, S3R), highlighting FRMD8’s role in tmTNF-α regulation. These data suggest that FRMD8 knockdown inhibits TNF-α secretion and increases tmTNF-α expression, acting through cell contact rather than soluble mediators.

Since sTNF-α binds TNFR1 and tmTNF-α binds both TNFR1 and TNFR2, we hypothesized that FRMD8 knockdown enhances TNFR2 signaling. In the GSE161529 dataset, low FRMD8 expression increased TNF-TNFRSF1B signaling in BRCA1 mutant tumors, enhancing TNFR2 intensity (Figure 3O). Low-FRMD8 tissues showed increased TNF signaling between tumor and immune cells (Figure S3S) and elevated TNF signaling strength (Figure S3T). Thus, FRMD8 knockdown likely enhances TNFR2’s role, supported by increased tmTNF-α. Treating cells with the TNFR1 inhibitor R-7050 and the TNFR1/TNFR2 inhibitor Etanercept showed reduced EMT markers and proliferation, especially with TNFR1/TNFR2 inhibition (Figures 3P, 3Q), indicating that EMT pathway activation and proliferation depend more on TNFR2 with FRMD8 knockdown, correlating with increased tmTNF-α.

### FRMD8 Directly Targets iRhom2 Protein and Promotes tmTNF-**α** Expression

Existing literature indicates that FRMD8 regulates iRhom2 stability, affecting ADAM17 maturation and cleavage, and subsequent TNF-α shedding. Our data show that FRMD8 knockdown inhibits iRhom2 expression and ADAM17 maturation (Figure 4A), consistent across MCF-7, BT-549, and MDA-MB-231 cells (Figure S4A). Flow cytometry revealed that FRMD8 knockdown reduced membrane iRhom2 expression (Figures 4B, 4C, S4B–S4G), without affecting ADAM17 levels (Figures 4D, 4E). Immunofluorescence confirmed reduced iRhom2 and E-cadherin colocalization on the membrane (Figure 4F), while ADAM17 levels remained unchanged (Figure 4G).

**Figure 4.**
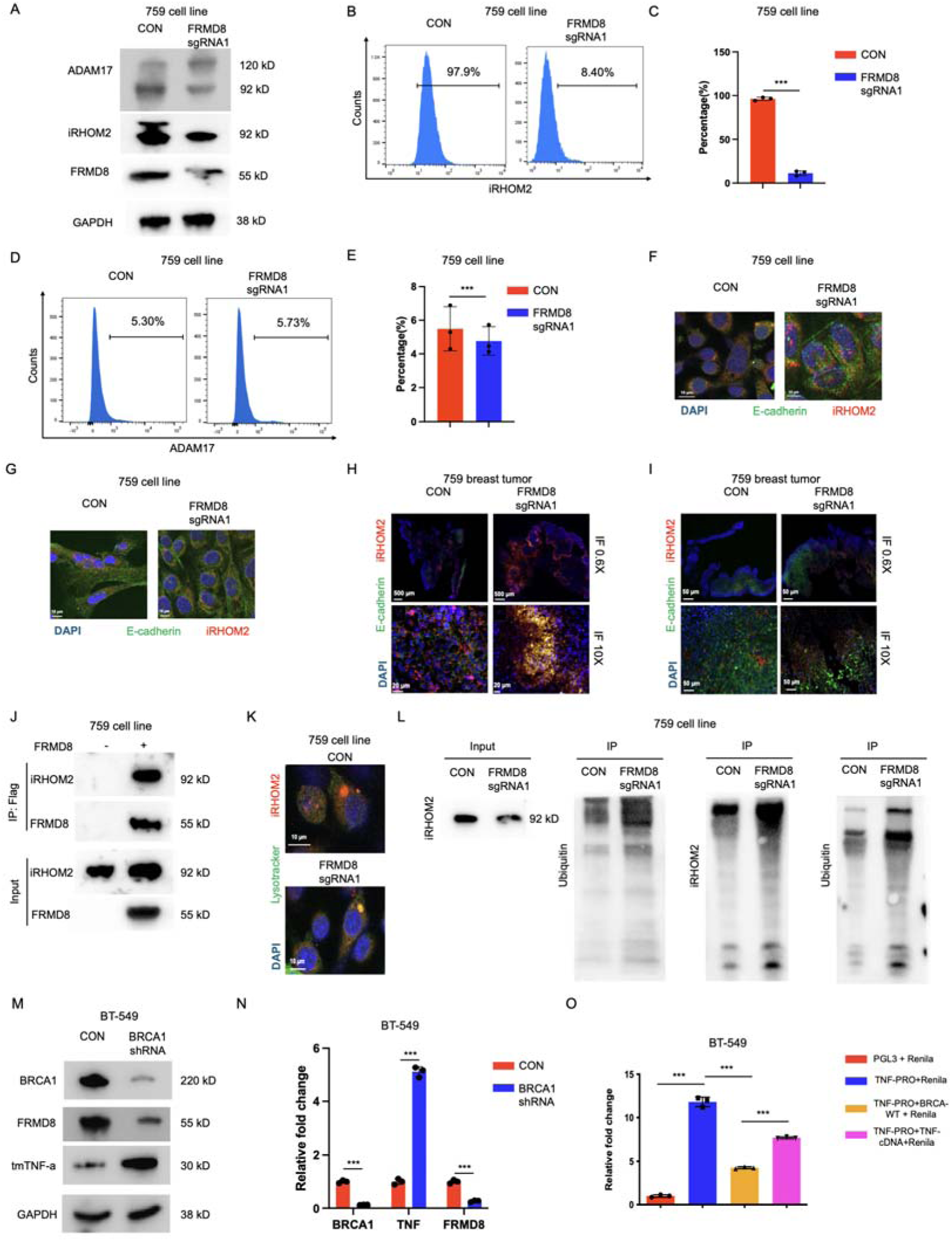
FRMD8 Directly Targets iRhom2 Protein and Promotes tmTNF-α Expression. (A) Western blot analysis of ADAM17, iRhom2, and FRMD8 protein expression in 759 cells transfected with FRMD8 sgRNA1, showing decreased iRhom2 and mature ADAM17 levels upon FRMD8 knockdown. (B, C) Flow cytometry analysis (B) and statistical analysis (C) of surface iRhom2 expression in 759 CON and FRMD8 sgRNA1 cells, indicating reduced membrane iRhom2 with FRMD8 knockdown. (D, E) Flow cytometry analysis (D) and statistical analysis (E) of surface ADAM17 expression, showing no significant changes. (F, G) Immunofluorescence staining of iRhom2 (F) and ADAM17 (G) in 759 CON and FRMD8 sgRNA1 cells, confirming decreased membrane iRhom2 without affecting ADAM17. (H, I) Immunofluorescence staining of iRhom2 (H) and ADAM17 (I) in primary tumors from 759 CON and FRMD8 sgRNA1 groups, supporting in vitro findings. (J) Co-immunoprecipitation assay demonstrating the interaction between FRMD8 and iRhom2 in 759 cells. (K) Immunofluorescence staining showing increased colocalization of iRhom2 with lysosomal marker LAMP1 in FRMD8 sgRNA1 cells, suggesting lysosomal degradation of iRhom2 upon FRMD8 knockdown. (L) Western blot analysis of ubiquitinated iRhom2 in 759 CON and FRMD8 sgRNA1 cells, indicating increased ubiquitination with FRMD8 knockdown. (M) Western blot analysis of BRCA1, FRMD8, and tmTNF-α in BT-549 cells transfected with BRCA1 shRNA, showing decreased BRCA1 and FRMD8 levels with increased tmTNF-α. (N) qPCR analysis of BRCA1, TNF-α, and FRMD8 expression in BT-549 CON and BRCA1 shRNA cells, confirming transcriptional changes. (O) Dual-luciferase reporter assay demonstrating the binding of BRCA1 to the TNF-α promoter, enhancing TNF-α transcription.Data are presented as mean ± SD. Statistical significance was assessed using a two-tailed Student’s t-test. ***P < 0.001.

In primary tumors from 759 CON and FRMD8 sgRNA1 mice, FRMD8 regulated iRhom2 membrane expression (Figure 4H) without significantly affecting ADAM17 (Figure 4I). mRNA analysis showed no changes in iRhom2 or ADAM17 transcription (Figure S4H).

Previous studies identified FRMD8’s role in stabilizing iRhom2 and promoting ADAM17 maturation, supported by our findings. Immunoprecipitation confirmed FRMD8’s interaction with iRhom2 (Figures 4J, S4I) and ADAM17 (Figures S4J, S4K). Immunofluorescence and immunoprecipitation indicated increased iRhom2 lysosomal localization and ubiquitin-bound ADAM17 upon FRMD8 knockdown (Figures 4K, 4L).

In BT-549 cells, BRCA1 overexpression reduced TNF-α and increased FRMD8 protein levels (Figure 4M) and mRNA levels (Figure 4N). BRCA1 was found to bind directly to the TNF promoter (Figure 4O). Furthermore, inhibition of BRCA1 in BT-549 and MDA-MB-231 cells greatly increased tmTNF-α on the cell membrane, whereas elevated BRCA1 decreased tmTNF-α (Figure S4L). Combining BRCA1, FRMD8, iRhom2, and ADAM17, we observed enrichment of TNF production and response pathways; however, FRMD8, iRhom2, and ADAM17 alone did not enrich the TNF pathway (Figure S4M), suggesting that BRCA1 modulates TNF transcription, maintaining high TNF levels in BRCA1-mutant cells.

### Low Expression of FRMD8/iRhom2 Significantly Promotes Tumor Metastasis In Vivo

Having established FRMD8’s regulation of iRhom2, we divided 759 cell lines into CON, FRMD8 sgRNA1, FRMD8 sgRNA1 + iRhom2 shRNA, and FRMD8 overexpression (OE) groups, each transfected with the respective vectors. We assessed the expression of ADAM17, iRhom2, FRMD8, and tmTNF-α, finding the lowest levels of FRMD8, iRhom2, and ADAM17, but the highest tmTNF-α in the FRMD8 sgRNA1 + iRhom2 shRNA group (Figure 5A). Immunofluorescence (Figure 5B) and flow cytometry (Figures 5C, 5D) confirmed the highest tmTNF-α levels in this group, suggesting the strongest metastatic potential.

**Figure 5.**
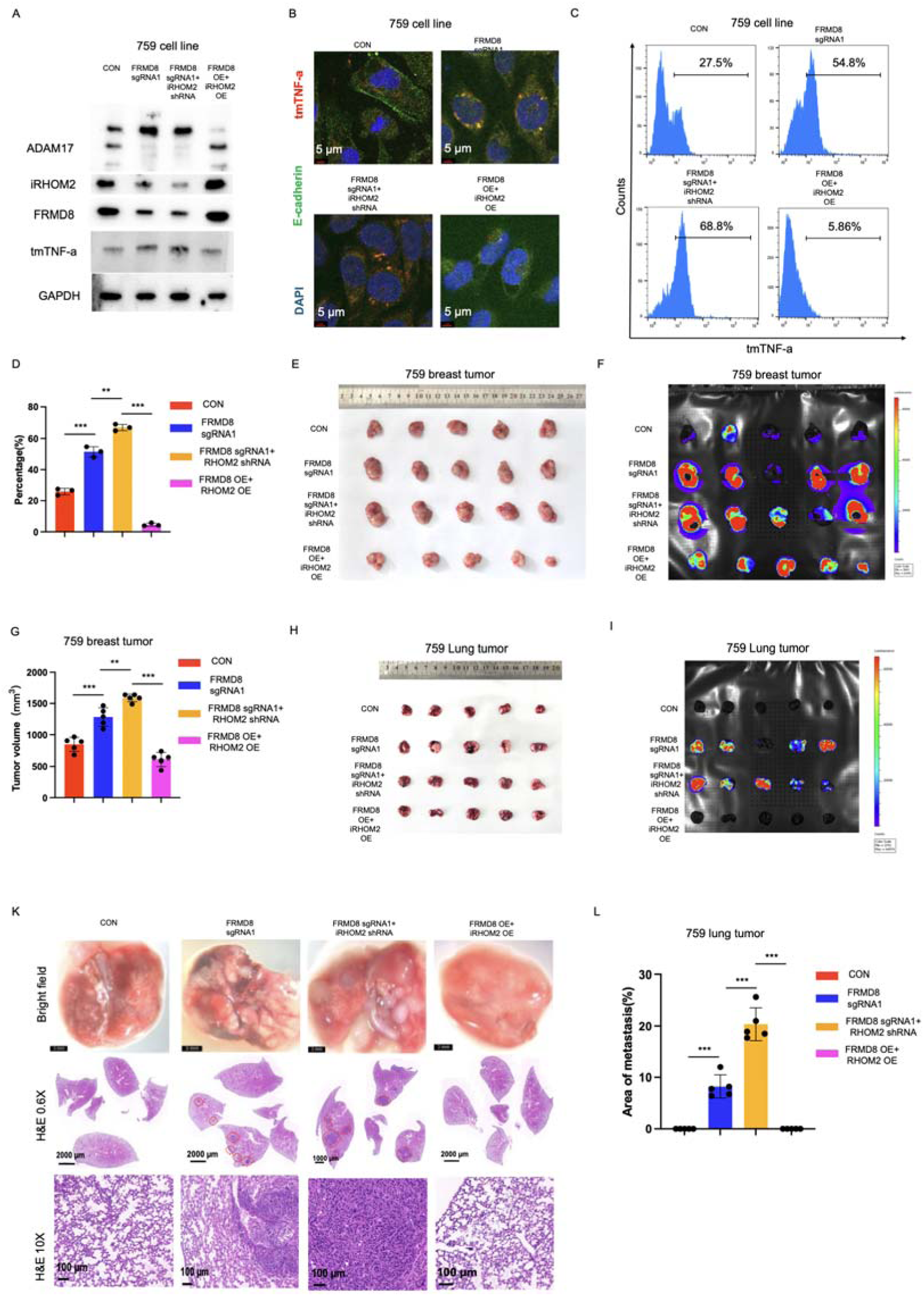
Low Expression of FRMD8/iRhom2 Significantly Promotes Tumor Metastasis In Vivo. (A) Western blot analysis of ADAM17, iRhom2, FRMD8, and tmTNF-α in 759 cells transfected with CON, FRMD8 sgRNA1, FRMD8 sgRNA1 + iRhom2 shRNA, and FRMD8 overexpression (OE), showing the lowest levels of FRMD8, iRhom2, and ADAM17 with the highest tmTNF-α in the FRMD8 sgRNA1 + iRhom2 shRNA group. (B) Immunofluorescence staining of tmTNF-α in the cells from (A), confirming increased tmTNF-α expression in the FRMD8 sgRNA1 + iRhom2 shRNA group. (C, D) Flow cytometry analysis (C) and statistical analysis (D) of surface tmTNF-α expression, supporting immunofluorescence results. (E) Images of primary tumors from mice injected with 759 CON, FRMD8 sgRNA1, FRMD8 sgRNA1 + iRhom2 shRNA, and FRMD8 OE cells, showing the largest tumors in the FRMD8 sgRNA1 + iRhom2 shRNA group. (F) In vivo imaging of GFP signals in primary tumors, indicating increased tumor growth in the FRMD8 sgRNA1 + iRhom2 shRNA group. (G) Statistical analysis of primary tumor volumes, confirming significant increases in the FRMD8 sgRNA1 + iRhom2 shRNA group. (H) Gross images of lungs from mice in each group, revealing visible lung metastases in the FRMD8 sgRNA1 + iRhom2 shRNA group. (I) Quantification of GFP signal intensity in lungs, indicating enhanced metastasis in the FRMD8 sgRNA1 + iRhom2 shRNA group. (K) H&E staining of lung tissues, showing increased metastasis in the FRMD8 sgRNA1 + iRhom2 shRNA group. (L) Statistical analysis of lung metastasis incidence, confirming significant increases in the FRMD8 sgRNA1 + iRhom2 shRNA group. Data are presented as mean ± SD (n = 5 mice per group). Statistical significance was assessed using a two-tailed Student’s t-test. ***P < 0.001.

We injected 759 CON, FRMD8 sgRNA1, FRMD8 sgRNA1 + iRhom2 shRNA, and FRMD8 OE cells into nude mice. The FRMD8 sgRNA1 + iRhom2 shRNA group showed the largest tumor volumes (Figures 5E, 5G) and the strongest GFP signals (Figure 5F). Visible lung metastases were observed in this group (Figure 5H), with GFP intensity significantly higher than the FRMD8 sgRNA1 group (Figure 5I). The highest lung metastasis levels were found in the FRMD8 sgRNA1 + iRhom2 shRNA group (Figures 5K, 5L). This group also exhibited the highest metastatic signals in the liver (Figures S5A, S5B), spleen (Figures S5C, S5D), brain (Figures S5E, S5F), and kidneys (Figures S5G, S5H). These data indicate that FRMD8 sgRNA1 + iRhom2 shRNA significantly enhances tmTNF-α expression and promotes in vivo tumor proliferation and metastasis.

### Knockdown of FRMD8 Recruits MDSCs in Primary Tumors and Enhances Their Function

Given that FRMD8 knockdown promotes CCL3 secretion, which is linked to immune cell recruitment, we employed CyTOF to comprehensively profile immune cells in CON and FRMD8 sgRNA1 groups (Figures 6A, S6A). FRMD8 knockdown increased the proportions of granulocytic MDSCs (G-MDSCs) and tumor cells (Figure 6B). Flow cytometry confirmed elevated G-MDSC levels post-FRMD8 knockdown (Figures 6C, 6D), and an increase in monocytic MDSCs (M-MDSCs) was also observed (Figures S6B, S6C). Elevated Gr1⁺ cell proportions further supported MDSC recruitment (Figure 6E).

**Figure 6.**
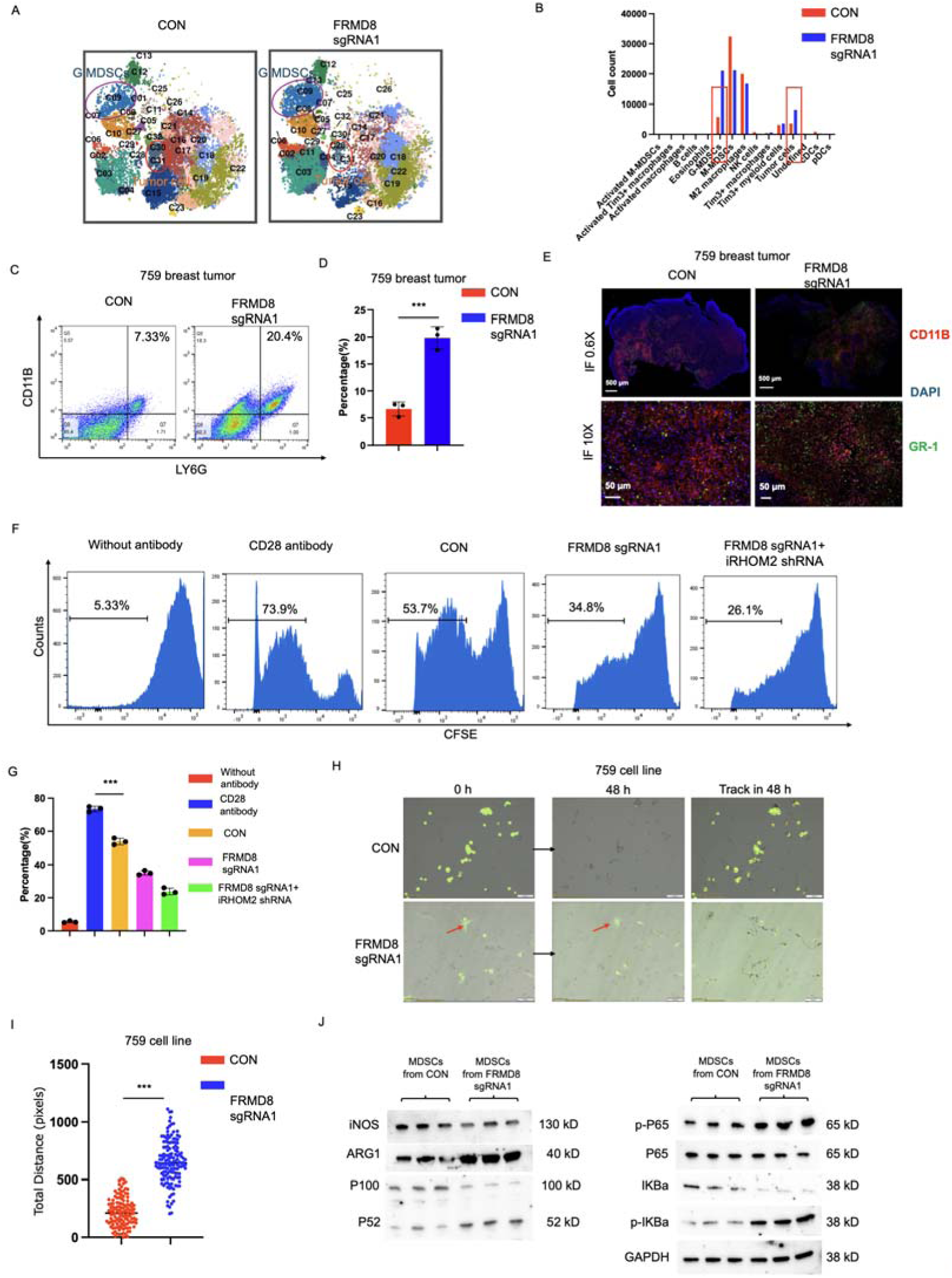
Knockdown of FRMD8 Recruits MDSCs to Primary Tumors and Enhances Their Function. (A) CyTOF analysis of immune cell subpopulations in primary tumors from 759 CON and FRMD8 sgRNA1 groups, illustrating increased proportions of G-MDSCs and tumor cells upon FRMD8 knockdown. (B) Statistical analysis highlighting the significant increase in G-MDSCs in the FRMD8 sgRNA1 group. (C, D) Flow cytometry analysis (C) and statistical analysis (D) of G-MDSC content in primary tumors, confirming CyTOF results. (E) Immunofluorescence staining of Gr1⁺ cells (MDSCs) in primary tumors, showing elevated MDSC infiltration with FRMD8 knockdown. (F) T cell suppression assay where MDSCs isolated from primary tumors of CON, FRMD8 sgRNA1, and FRMD8 sgRNA1 + iRhom2 shRNA groups were co-cultured with spleen-derived T cells, demonstrating increased T cell suppression by MDSCs from FRMD8 knockdown groups. (G) Statistical analysis of T cell proliferation, indicating significant inhibition by MDSCs from FRMD8 sgRNA1 and FRMD8 sgRNA1 + iRhom2 shRNA groups. (H) Co-culture of 759 CON and FRMD8 sgRNA1 cells with MDSCs, showing enhanced cell migration and interaction in the FRMD8 sgRNA1 group after 48 hours. (I) Statistical analysis of cell migration speed, confirming increased migration in the FRMD8 sgRNA1 group. (J) Western blot analysis of iNOS, ARG1, and NF-κB pathway activation in MDSCs isolated from primary tumors, indicating enhanced MDSC activation upon FRMD8 knockdown. Data are presented as mean ± SD. Statistical significance was assessed using a two-tailed Student’s t-test. ***P < 0.001.

We isolated Gr1⁺ cells from 759 CON, FRMD8 sgRNA1, and FRMD8 sgRNA1 + iRhom2 shRNA primary tumors and co-cultured them with T cells from the spleen (Figures 6F, 6G) and bone marrow (Figures S6D, S6E). Compared to the CON group, FRMD8 sgRNA1 significantly inhibited T cell proliferation, with FRMD8 sgRNA1 + iRhom2 shRNA showing more pronounced effects. Co-culture of CON and FRMD8 sgRNA1 tumor cells with MDSCs revealed that FRMD8 sgRNA1 tumor cells coexisted with MDSCs, enhancing cell migration distances (Figures 6H, 6I, S6F–S6I).

Lastly, Gr1⁺ cells isolated from 759 CON and FRMD8 sgRNA1 primary tumors displayed activation of both classical and non-classical NF-κB pathways, with significantly increased ARG1 expression, a marker of MDSC activation (Figure 6J). This indicates that arginase-related immune suppression pathways are significantly enhanced in MDSCs, consistent with the CyTOF results.

### Paclitaxel Inhibits FRMD8-Related Metastasis by Reversing FRMD8 Expression

We sought drugs that could block FRMD8-related metastasis. Analysis of the TCGA dataset revealed that patients responsive to paclitaxel treatment showed significantly increased FRMD8 expression (Figure 7A). Additionally, iRhom2 (Figure S7A) and ADAM17 (Figure S7B) levels were elevated, which was confirmed in the GSE22513 dataset (Figures S7C–S7E). Although anastrozole and doxorubicin showed similar effects in TCGA data, only paclitaxel increased both FRMD8 and iRhom2 expression in 759 cells (Figures 7B, 7C). Docetaxel, a paclitaxel analogue, yielded similar results (Figure S7F).

**Figure 7.**
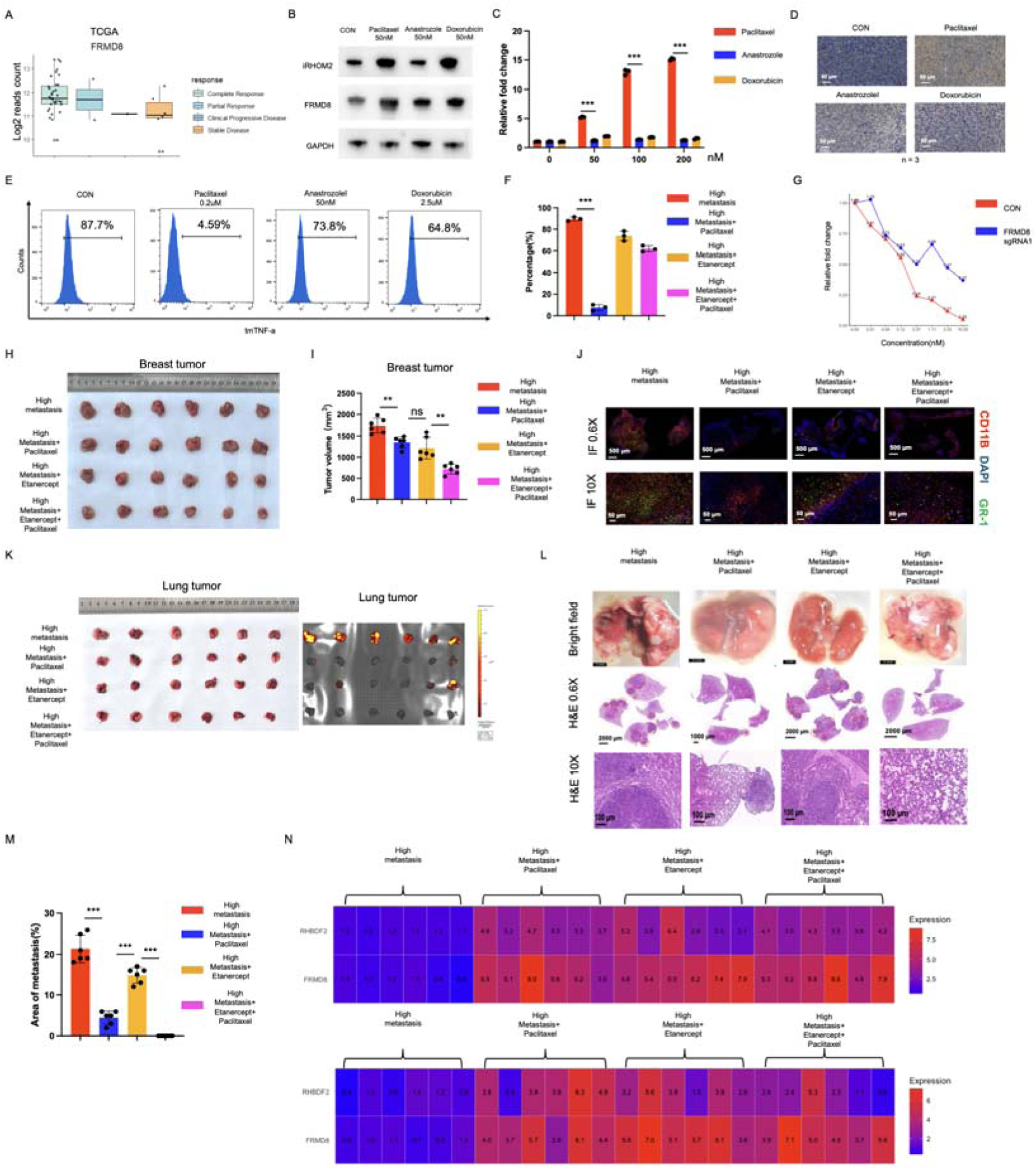
Paclitaxel Inhibits FRMD8-Related Metastasis by Reversing FRMD8 Expression. (A) Analysis of FRMD8 expression in paclitaxel responders and non-responders in the TCGA dataset, showing higher FRMD8 levels in responders. (B) Western blot analysis of iRhom2 and FRMD8 protein levels in 759 cells treated with paclitaxel, anastrozole, and doxorubicin for 24 hours, indicating increased FRMD8 and iRhom2 with paclitaxel treatment. (C) qPCR analysis of FRMD8 mRNA levels under the same treatments, supporting protein level findings. (D) Immunohistochemical staining of FRMD8 in primary tumors from mice injected with 759 FRMD8 sgRNA1 + iRhom2 shRNA cells and treated with paclitaxel, anastrozole, or doxorubicin, showing increased FRMD8 expression with paclitaxel. (E, F) Flow cytometry analysis (E) and statistical analysis (F) of surface tmTNF-α in 759 cells after treatments, demonstrating reduced tmTNF-α with paclitaxel. (G) MTT assay assessing drug sensitivity of 759 CON and 759 FRMD8 sgRNA1 cells to paclitaxel, showing increased sensitivity in FRMD8 knockdown cells. (H, I) Images (H) and statistical analysis (I) of primary tumor volumes in mice injected with 759 FRMD8 sgRNA1 + iRhom2 shRNA cells and treated with paclitaxel, Etanercept, or both, indicating significant tumor reduction with combined treatment. (J) Immunofluorescence staining of Gr1⁺ cells in primary tumors, showing decreased MDSC infiltration with treatments. (K–M) Gross images (K), H&E staining (L), and statistical analysis (M) of lung metastasis, demonstrating effective inhibition of metastasis with combined paclitaxel and Etanercept treatment. (N) qPCR analysis of FRMD8 and RHBDF2 (iRhom2) expression in breast and lung tumors, confirming increased expression with treatments. Data are presented as mean ± SD (n = 5 mice per group). Statistical significance was assessed using a two-tailed Student’s t-test. NS, not significant; **P < 0.01; ***P < 0.001.

Paclitaxel treatment increased FRMD8 expression in vivo (Figure 7D) and significantly reduced membrane tmTNF-α levels in vitro (Figures 7E, 7F). To comprehensively screen for drugs targeting low FRMD8 expression, we used a library of 146 FDA-approved drugs. FRMD8 sgRNA1 cells were highly sensitive to paclitaxel (Figure 7G) and docetaxel (Figure S7G), showing the highest sensitivity among the drugs tested (Figure S7H).

We defined 759 FRMD8 sgRNA1 + iRhom2 shRNA as the high-metastasis group, injected these cells into nude mice, and treated them with paclitaxel, Etanercept, or both. Paclitaxel alone reduced primary tumor volume and GFP signal intensity, as did Etanercept. Combined treatment significantly decreased tumor volume and GFP signal (Figures 7H, 7I, S7I). Paclitaxel and Etanercept co-treatment also reduced MDSC infiltration in primary tumors (Figure 7J) and blocked lung metastasis (Figures 7K–7M), kidney, liver, and brain metastasis (Figures 7N–7P), although no visible spleen metastasis was observed (Figure S7J).

qPCR confirmed that paclitaxel, Etanercept, and their combination increased FRMD8 and RHBDF2 (iRhom2) expression in breast and lung tumors, indicating that these treatments reversed FRMD8 and iRhom2 expression, thus blocking FRMD8/iRhom2-related metastasis.

## Discussion

Library screening is an efficient method for identifying target genes involved in disease processes [13]. In previous studies, we identified ATP11B as a metastasis-related gene using a whole-genome knockout library. In the current study, we combined sequencing data from our hospital with public datasets to identify 41 candidate genes associated with breast cancer metastasis. Of these, 39 genes were found to be present in mice. Screening these genes in the 759 cell line led to the identification of FRMD8 as a novel breast cancer metastasis-related gene.

Our analysis revealed that low expression of FRMD8 significantly increased the levels of tmTNF-α, with iRhom2 identified as a direct target. The reduced expression of FRMD8 and iRhom2 further promoted tmTNF-α expression and tumor metastasis, enhancing immune suppression pathways within the tumor microenvironment. These findings align with previous studies highlighting the role of tmTNF-α in promoting tumor progression and metastasis through activation of NF-κB pathways and modulation of the tumor microenvironment [14–17].

Existing literature highlights various functions of FRMD8, including the stabilization of the iRhom/ADAM17 complex, regulation of Wnt signaling, maintenance of cellular stability, and tumor suppression [18–20]. Our study confirmed FRMD8’s role in stabilizing membrane proteins in breast cancer. Despite the potential for cell cycle regulation pathways, no direct enrichment was observed in our analysis. Knockdown of FRMD8 resulted in decreased TNF secretion while increasing tmTNF-α expression. This suggests that inhibition of FRMD8 blocks the cleavage of tmTNF-α into soluble TNF-α, thereby promoting tmTNF-α expression and its associated pro-metastatic effects.

FRMD8 regulates the lysosomal degradation of iRhom2, which influences the cleavage and activity of ADAM17. This, in turn, affects downstream molecules such as TNF-α and EGFR [21–23]. Our study demonstrated that FRMD8 interacts with both iRhom2 and ADAM17, regulating iRhom2 expression on the cell membrane without altering ADAM17 membrane levels. Reduced expression of FRMD8 and iRhom2 led to increased tmTNF-α expression and tumor metastasis.

BRCA1 is primarily known for its role in DNA repair, but recent studies have demonstrated that BRCA1 also regulates inflammatory and immune responses, including the transcription and function of TNF-α [24]. BRCA1 can bind to the TNF promoter, enhancing TNF transcription levels [25–27]. In our research, we identified that BRCA1 deficiency promotes the membrane expression of TNF-α. The interaction among BRCA1, FRMD8, iRhom2, and ADAM17 facilitates TNF production and the activation of the TNF signaling pathway. These findings suggest that in BRCA1-deficient tissues, the TNF-α pathway is further activated.

Unlike sTNF-α, which primarily activates the classical NF-κB pathway via TNFR1, tmTNF-α can bind both TNFR1 and TNFR2, activating both classical and non-classical NF-κB pathways. This leads to the expression of metastasis-related molecules, and tmTNF-α also plays a crucial role in activating MDSCs and regulatory T cells in the tumor microenvironment, thereby enhancing immune suppression pathways [28]. Our study demonstrated increased TNFR2 signaling in FRMD8 low-expression tumors, confirming that FRMD8’s function is heavily dependent on tmTNF-α.

In vitro experiments demonstrated that tmTNF-α promotes the activation of the EMT pathway, a crucial process in cancer metastasis. In vivo, tmTNF-α enhances tumor progression by modulating immune responses. FRMD8 knockdown resulted in increased expression of CCL3, a known inducer of MDSCs. CyTOF screening confirmed elevated levels of MDSCs in tissues with FRMD8 knockdown. Further studies showed that both classical and non-classical NF-κB pathways were activated in MDSCs, reinforcing the role of FRMD8 in MDSC activation.

Combining drug library screening and public datasets, we identified paclitaxel as a compound capable of reversing the expression of FRMD8, iRhom2, and ADAM17. FRMD8 knockdown cells showed increased sensitivity to paclitaxel, suggesting that paclitaxel effectively blocks FRMD8-related metastasis by altering FRMD8 expression. In vivo experiments demonstrated that paclitaxel partially inhibited FRMD8-related metastasis. While Etanercept, a TNF inhibitor [29], partially reduced metastasis, it was not fully effective on its own. However, the combination of paclitaxel and Etanercept completely blocked FRMD8-related metastasis, indicating that this combination therapy could effectively prevent FRMD8-related metastasis and may serve as a promising therapeutic strategy for breast cancer patients.

Our study suggests that FRMD8-related metastasis involves multiple molecular mechanisms, including the regulation of tmTNF-α, activation of NF-κB pathways, and modulation of immune cell functions. The interaction between FRMD8 and iRhom2 plays a crucial role in maintaining ADAM17 stability and function, which is essential for tmTNF-α cleavage and subsequent activation of metastasis-related pathways. Additionally, the increased expression of CCL3 in FRMD8 knockdown cells and tissues indicates a significant role in recruiting and activating MDSCs, which further supports tumor growth and immune evasion.

The identification of FRMD8 as a key regulator of breast cancer metastasis has significant clinical implications. Targeting FRMD8 and its associated pathways could provide new therapeutic strategies for preventing and treating metastatic breast cancer. The combination of paclitaxel and Etanercept shows promise in effectively inhibiting FRMD8-related metastasis, suggesting potential for clinical application. Future research should focus on further elucidating the molecular mechanisms underlying FRMD8’s role in metastasis and exploring additional therapeutic agents that can target these pathways.

In conclusion, our study identified FRMD8 as a key player in breast cancer metastasis. Low expression of FRMD8 promotes tmTNF-α expression and enhances tumor metastasis through multiple mechanisms, including the activation of NF-κB pathways and immune suppression. The combination of paclitaxel and Etanercept shows potential as an effective therapy to inhibit FRMD8-related metastasis. Further research is warranted to explore the full therapeutic potential of targeting FRMD8 and its associated pathways in breast cancer treatment.

## Author Contributions

Conception and design: Haibo Wu

Development of methodology: Jun Xu

Acquisition of data (provided animals, acquired and managed patients, provided facilities, etc.):Xiaoyu Yang

Analysis and interpretation of data (e.g., statistical analysis, biostatistics, computational analysis): Jun XU

Writing, review, and/or revision of the manuscript: Zhe Wang

## Acknowledgments

The authors thank the members of the Wu and Wang laboratories for helpful advice and discussions. We also acknowledge the Animal Research Core for providing animal housing. This work was supported by This work was supported by the National Natural Science Foundation of China(82303451).

---

**Supplementary Figure 1.**
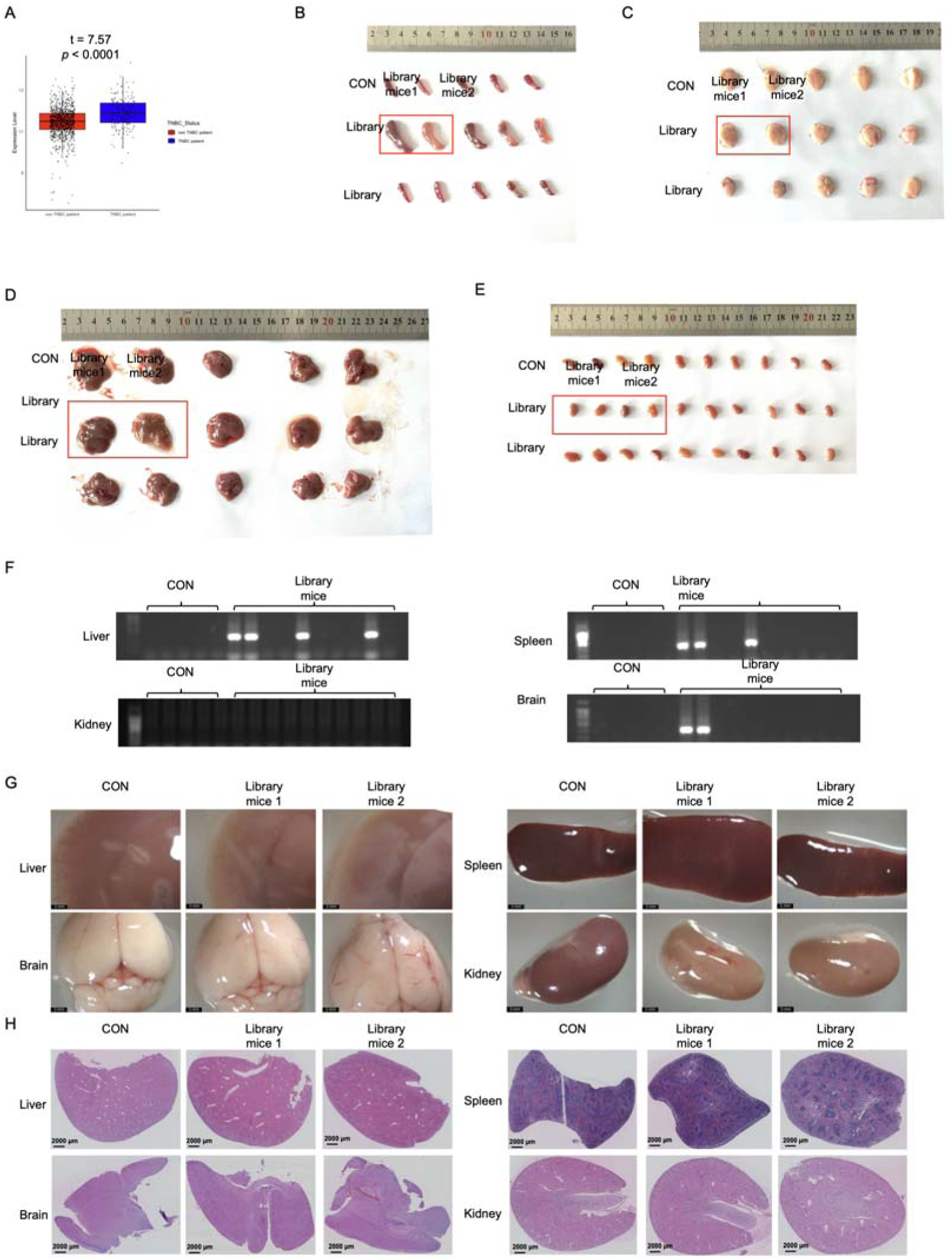
Additional Analysis Supporting FRMD8 as a Metastasis-Related Gene. (A) Analysis of FRMD8 expression levels in triple-negative breast cancer (TNBC) and non-TNBC tumors in the TCGA dataset, showing significantly lower FRMD8 expression in TNBC tumors. (B–E) Gross images of spleen (B), brain (C), liver (D), and kidneys (E) from control mice and library mice, illustrating organ changes in library mice. (F) PCR detection of metastasis in liver, spleen, brain, and kidneys of control mice and library mice, confirming the presence of FRMD8 sgRNA in multiple organs of library mice. (G–H) Microscopic images (G) and H&E staining (H) of liver, kidney, spleen, and brain tissues from control mouse 1, library mouse 1, and library mouse 2, showing histopathological changes in library mice.Data are presented as mean ± SD (n = 5 mice per group). Statistical significance was assessed using a two-tailed Student’s t-test. ***P < 0.001.

**Supplementary Figure 2.**
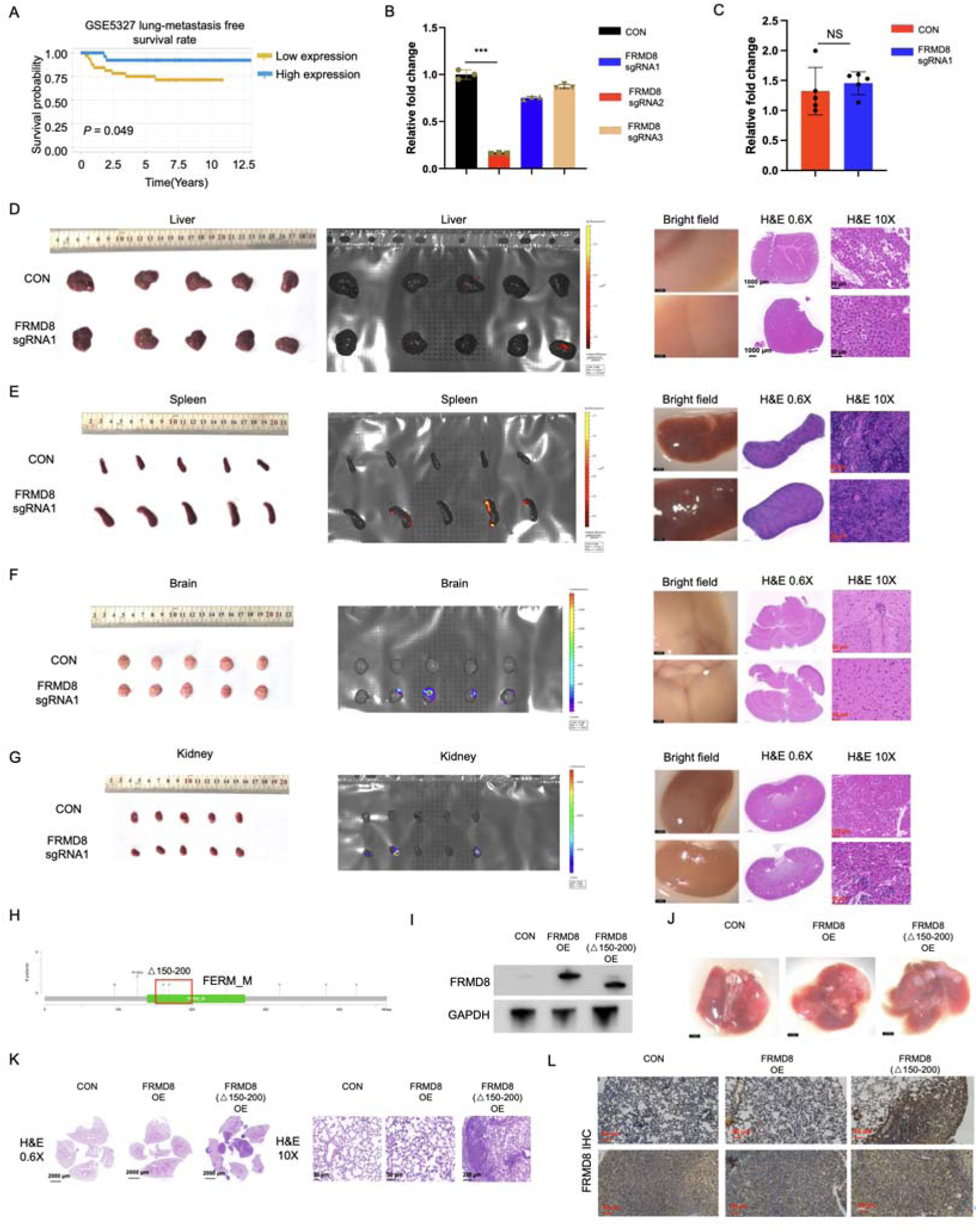
Additional Analysis of FRMD8’s Role in Metastasis. (A) Kaplan–Meier survival analysis showing the correlation between FRMD8 expression and lung metastasis survival rates in the GSE5327 dataset, indicating poorer survival with low FRMD8 expression. (B) Quantification of Western blot bands from Figure 2C, confirming significant FRMD8 knockdown by sgRNA1. (C) Statistical analysis of lung volumes in mice injected with 759 CON and 759 FRMD8 sgRNA1 cells, showing no significant difference between groups. (D–G) Gross images, GFP signal measurements, and H&E staining of liver (D), spleen (E), brain (F), and kidneys (G) from 759 CON and 759 FRMD8 sgRNA1 groups, indicating increased metastasis in the FRMD8 sgRNA1 group. (H) Structural diagram of the FRMD8 protein, highlighting the FERM domain and the Δ150–200 deletion region. (I) Western blot analysis of FRMD8 expression in 759 cells transfected with FRMD8 cDNA and FRMD8(Δ150–200), showing slightly reduced molecular weight in the deletion mutant. (J–L) Gross images, H&E staining, and IHC analysis of lung metastasis in mice injected with 759 CON, 759 FRMD8 OE, and 759 FRMD8(Δ150–200) OE cells, demonstrating increased metastasis with the FRMD8(Δ150–200) mutant. Data are presented as mean ± SD (n = 5 mice per group). Statistical significance was assessed using a two-tailed Student’s t-test. NS, not significant; ***P < 0.001.

**Supplementary Figure 3.**
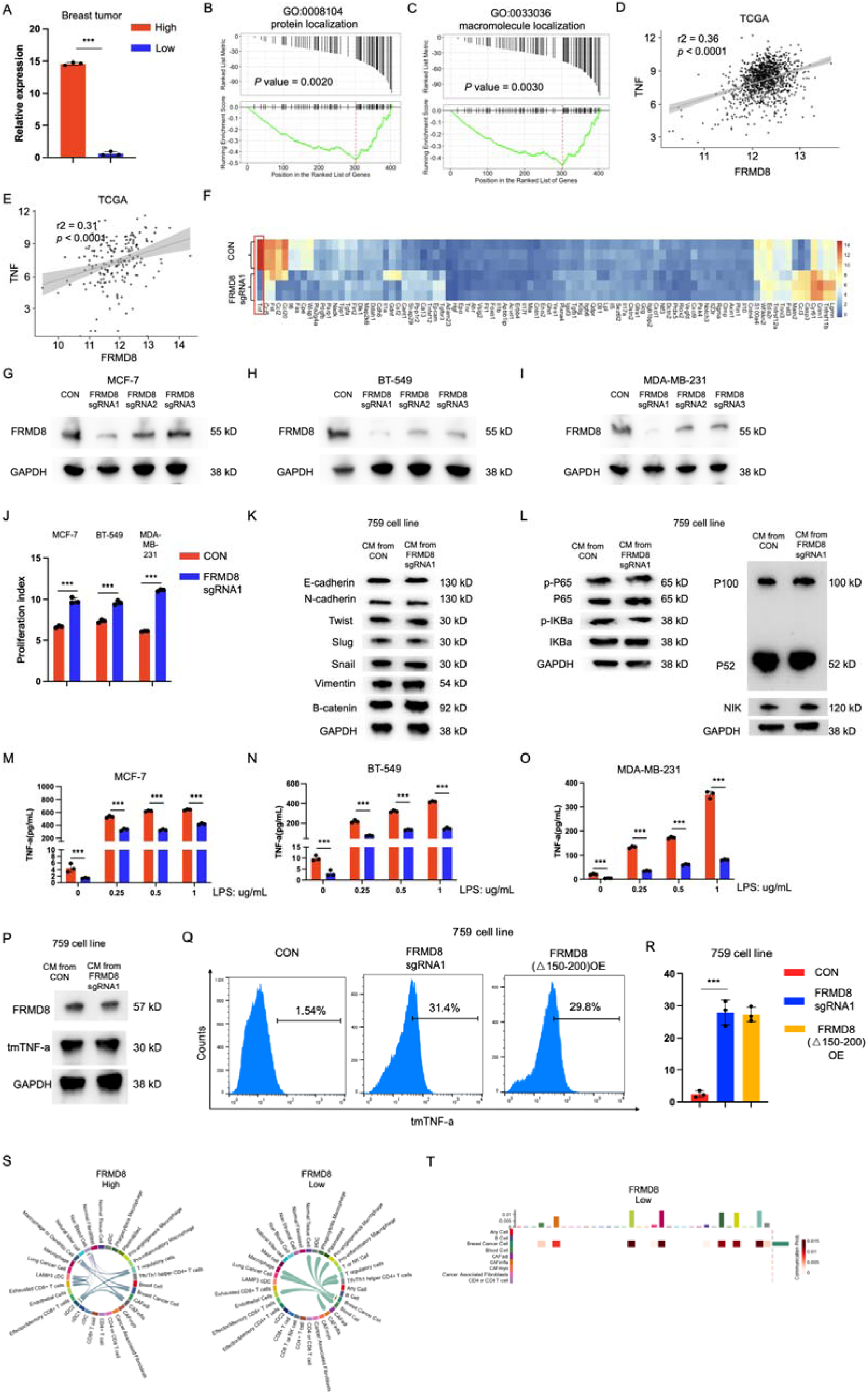
Additional Analysis of FRMD8 Knockdown Effects. (A) qPCR confirmation of FRMD8 expression levels before sequencing in tumors with low and high FRMD8 expression. (B, C) Pathway enrichment analysis in CON and FRMD8 sgRNA1 cells from our hospital’s dataset, supporting the findings in Figure 3A. (D, E) Pearson correlation analysis between FRMD8 and TNF-α expression in the TCGA and TNBC datasets, showing a positive correlation. (F) Cytokine array analysis highlighting increased CCL3 expression in the supernatant of FRMD8 sgRNA1 cells. (G–I) Western blot analysis of FRMD8 protein levels in MCF-7 (G), BT-549 (H), and MDA-MB-231 (I) cells transfected with CON, FRMD8 sgRNA1, FRMD8 sgRNA2, and FRMD8 sgRNA3, confirming effective knockdown by sgRNA1. (J) Cell proliferation assays in MCF-7, BT-549, and MDA-MB-231 CON and FRMD8 sgRNA1 cells, showing increased proliferation upon FRMD8 knockdown. (K) Western blot analysis of EMT markers in 759 cells treated with supernatants from 759 CON and 759 FRMD8 sgRNA1 cells, indicating no significant changes, supporting cell-autonomous effects. (L) Western blot analysis of NF-κB pathway activation in 759 cells treated with supernatants from 759 CON and 759 FRMD8 sgRNA1 cells, showing no significant activation. (M–O) ELISA measurements of TNF-α secretion in MCF-7 (M), BT-549 (N), and MDA-MB-231 (O) CON and FRMD8 sgRNA1 cells, confirming reduced TNF-α secretion upon FRMD8 knockdown. (P) Flow cytometry analysis of tmTNF-α in 759 cells treated with supernatants from 759 CON and 759 FRMD8 sgRNA1 cells, showing no significant changes. (Q, R) Flow cytometry analysis (Q) and statistical analysis (R) of tmTNF-α in 759 cells transfected with FRMD8 cDNA and FRMD8(Δ150–200), indicating increased tmTNF-α with the deletion mutant. (S) Analysis of TNF signaling intensity between tumor and immune cells in tissues with high and low FRMD8 expression in BRCA1 mutant tumors. (T) Analysis of TNF signaling inflow and outflow intensity in tissues with low FRMD8 expression in BRCA1 mutant tumors, supporting enhanced TNF signaling. Data are presented as mean ± SD. Statistical significance was assessed using a two-tailed Student’s t-test or Pearson correlation test. ***P < 0.001.

**Supplementary Figure 4.**
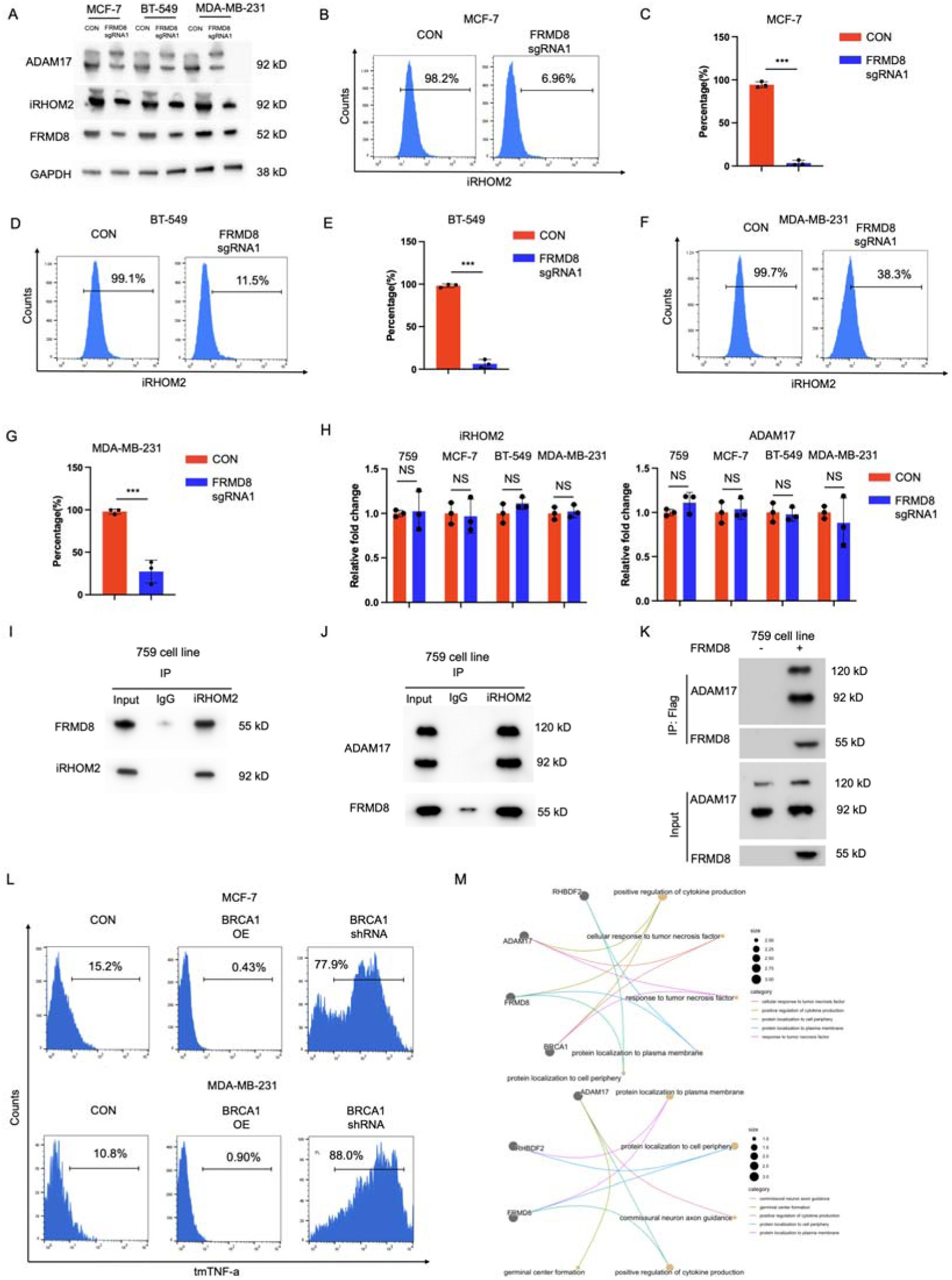
Additional Analysis of FRMD8 and iRhom2 Interaction. (A) Western blot analysis of ADAM17, iRhom2, and FRMD8 protein levels in MCF-7, BT-549, and MDA-MB-231 cells transfected with CON and FRMD8 sgRNA1, supporting findings in 759 cells. (B–G) Flow cytometry analysis and statistical analysis of surface iRhom2 expression in MCF-7 (B, C), BT-549 (D, E), and MDA-MB-231 (F, G) CON and FRMD8 sgRNA1 cells, confirming decreased iRhom2 expression upon FRMD8 knockdown. (H) qPCR analysis of iRhom2 and ADAM17 mRNA levels in 759, MCF-7, BT-549, and MDA-MB-231 CON and FRMD8 sgRNA1 cells, showing no significant transcriptional changes. (I–K) Co-immunoprecipitation assays demonstrating the interactions between FRMD8 and iRhom2 (I), and between FRMD8 and ADAM17 (J, K) in 759 cells. (L) Flow cytometry analysis of tmTNF-α in MCF-7 and MDA-MB-231 cells with BRCA1 overexpression or knockdown, showing that BRCA1 levels inversely affect tmTNF-α expression. (M) Gene set enrichment analysis in the TCGA-BRCA dataset investigating pathway alterations associated with BRCA1, FRMD8, ADAM17, and iRhom2, highlighting the impact on TNF-α signaling pathways. Data are presented as mean ± SD. Statistical significance was assessed using a two-tailed Student’s t-test. NS, not significant; ***P < 0.001.

**Supplementary Figure 5.**
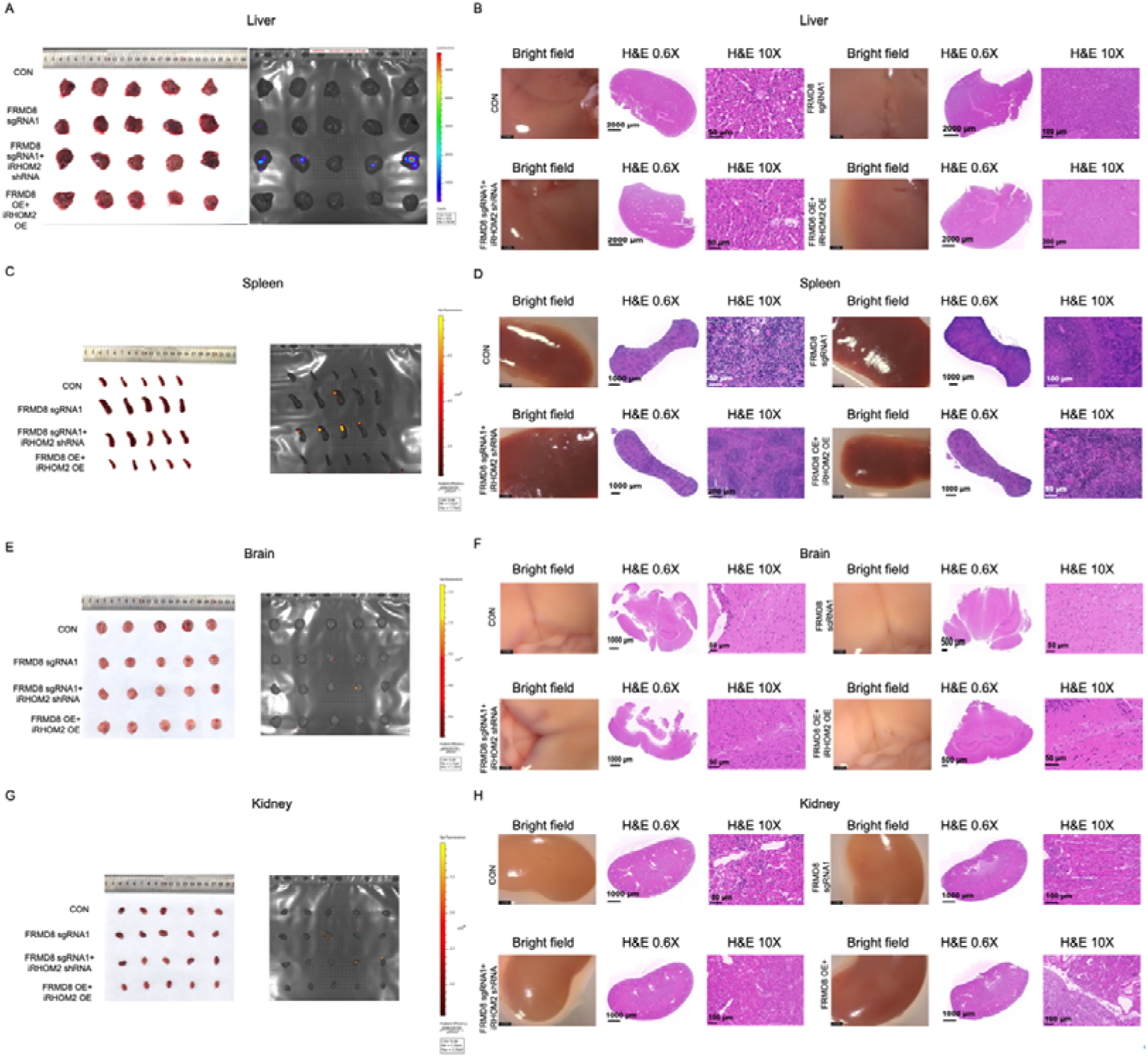
Additional Analysis of Metastasis in Other Organs. (A–B) Gross images, GFP signal measurements (A), and H&E staining (B) of liver tissues from mice injected with 759 CON, FRMD8 sgRNA1, FRMD8 sgRNA1 + iRhom2 shRNA, and FRMD8 OE cells, showing increased liver metastasis in the FRMD8 sgRNA1 + iRhom2 shRNA group. (C–D) Similar analysis for spleen tissues, indicating enhanced spleen metastasis. (E–F) Analysis of brain tissues, demonstrating increased brain metastasis. (G–H) Analysis of kidney tissues, confirming elevated kidney metastasis in the FRMD8 sgRNA1 + iRhom2 shRNA group. Data are presented as mean ± SD (n = 5 mice per group). Statistical significance was assessed using a two-tailed Student’s t-test. **P < 0.01; ***P < 0.001.

**Supplementary Figure 6.**
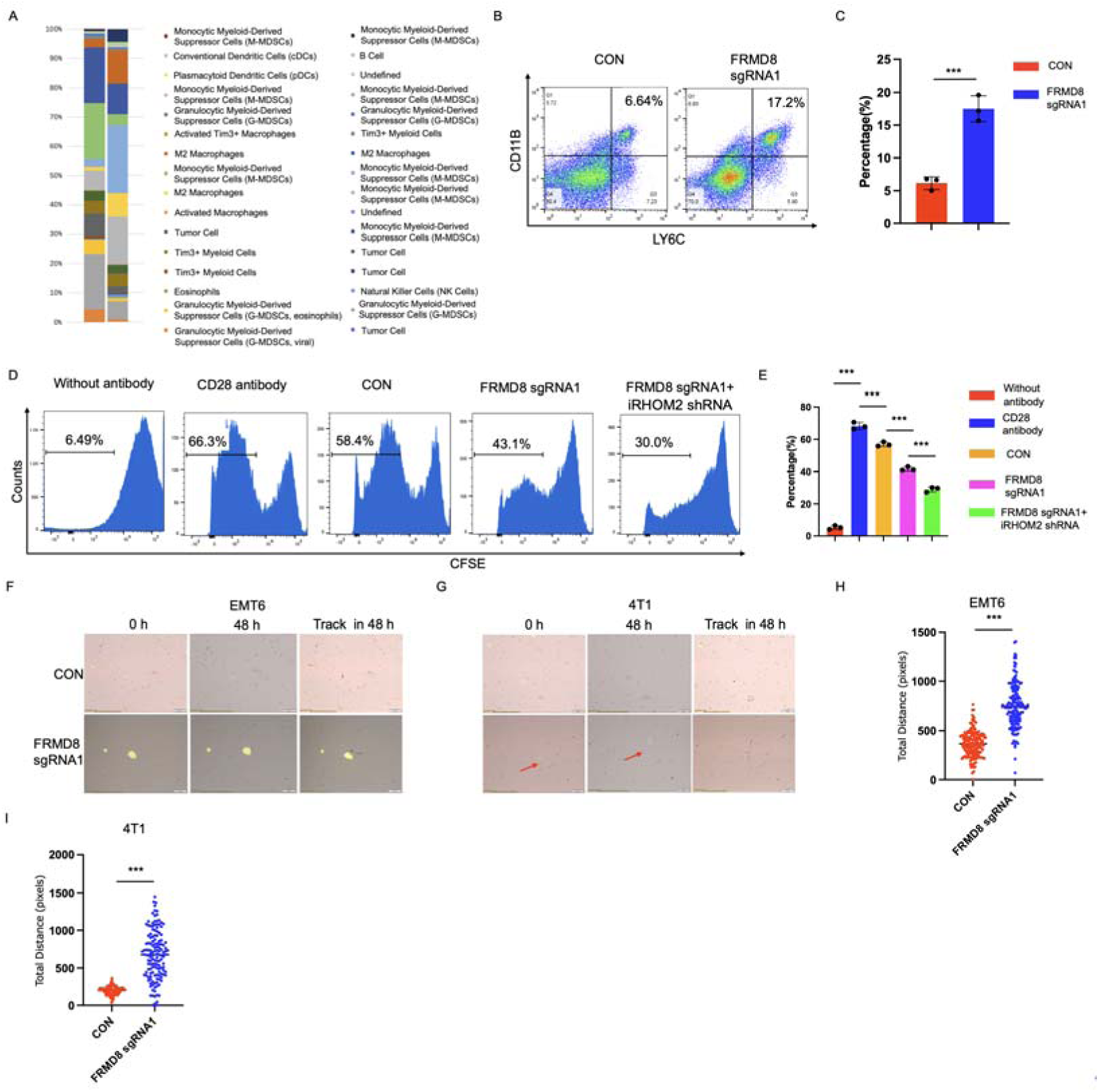
Additional Analysis of MDSC Recruitment and Function. (A) Annotation of immune cell populations in the CyTOF experiment. (B, C) Flow cytometry analysis (B) and statistical analysis (C) of M-MDSC proportions in primary tumors, showing increased M-MDSCs with FRMD8 knockdown. (D, E) T cell suppression assay using bone marrow-derived T cells, confirming enhanced T cell suppression by MDSCs from FRMD8 knockdown groups. (F–I) Co-culture experiments of EMT6 and 4T1 cell lines with MDSCs, demonstrating increased cell migration and interaction similar to findings with 759 cells. Data are presented as mean ± SD. Statistical significance was assessed using a two-tailed Student’s t-test. ***P < 0.001.

**Supplementary Figure 7.**
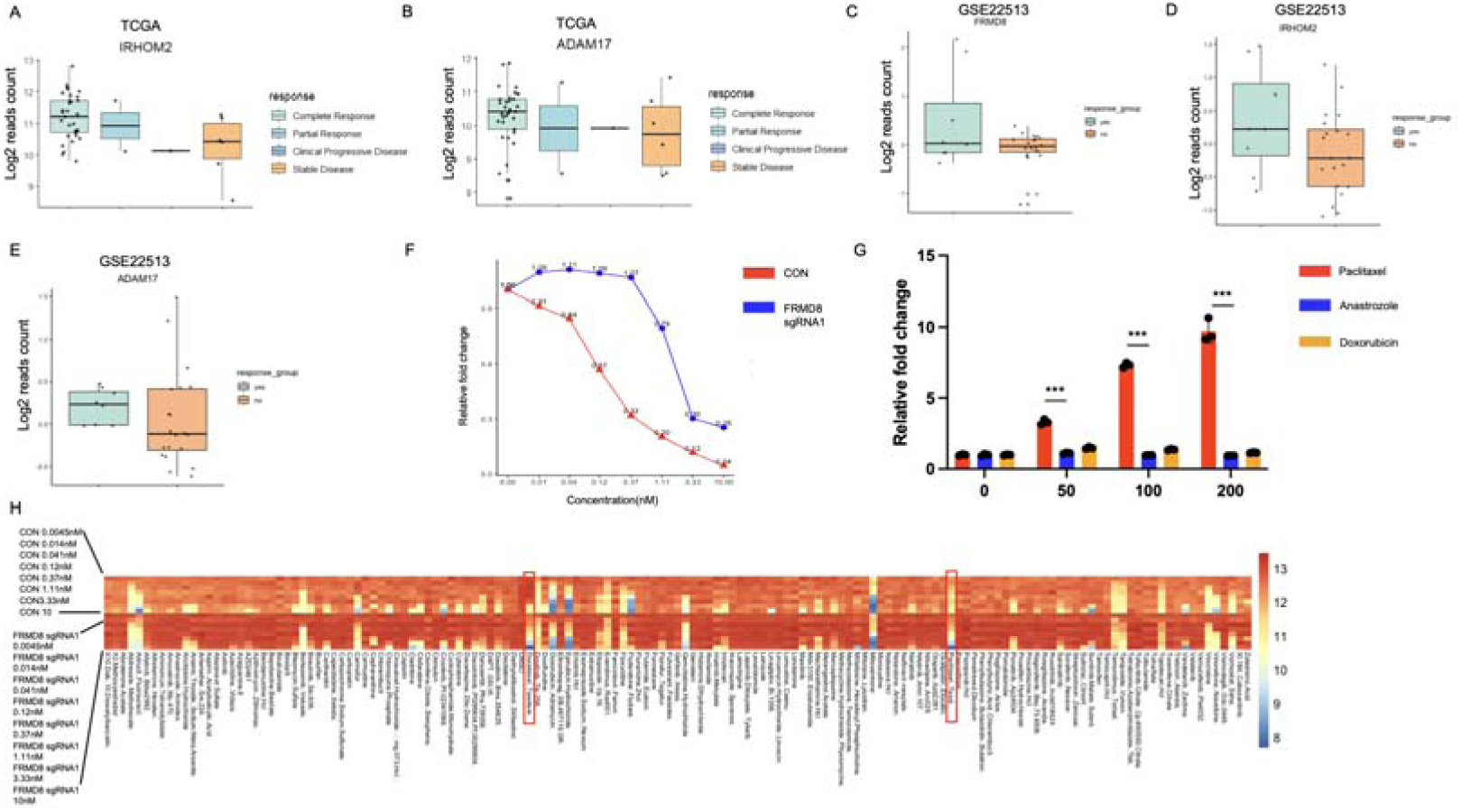
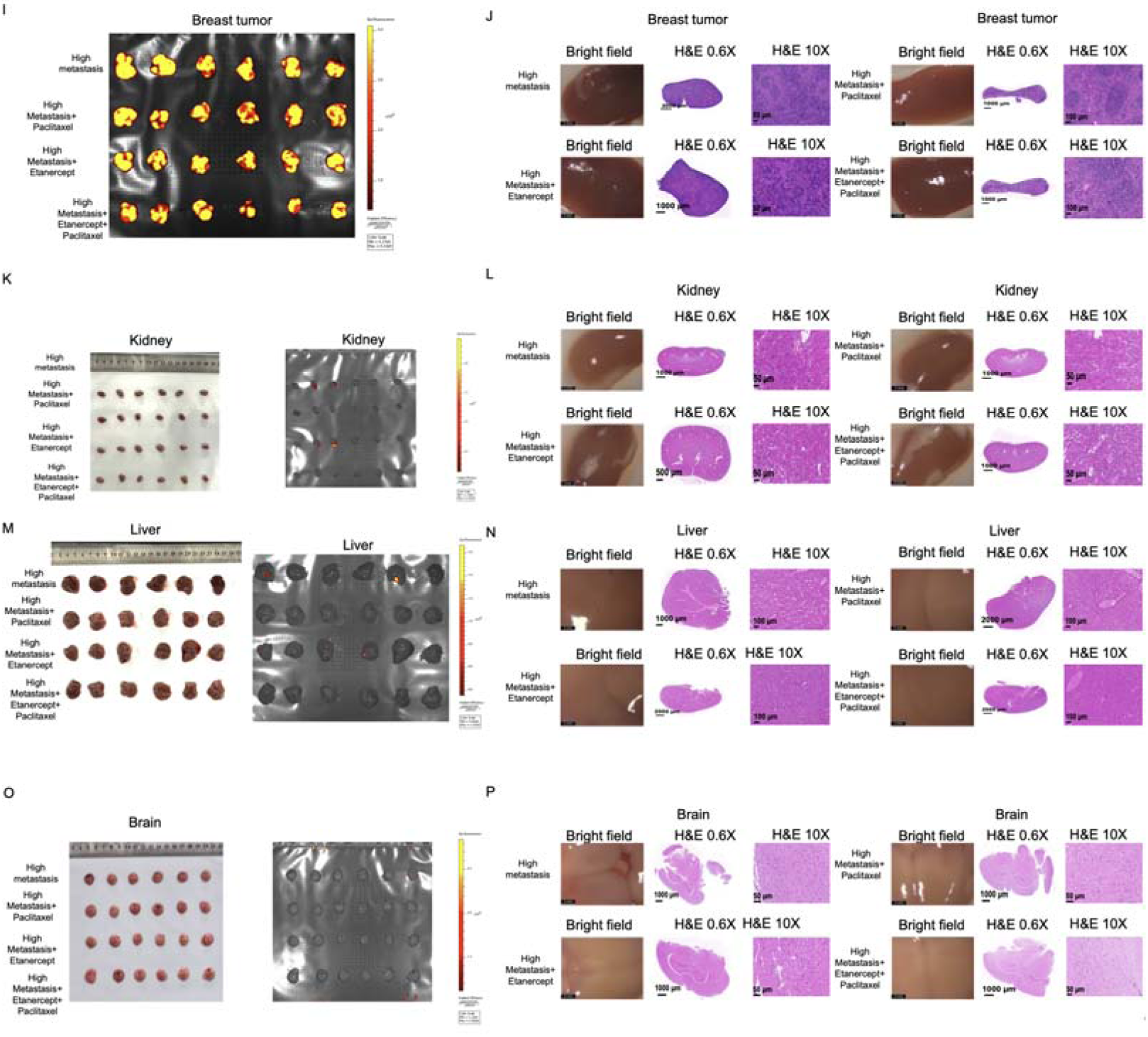
Additional Analysis of Drug Effects on FRMD8-Related Metastasis. (A, B) Analysis of iRhom2 (A) and ADAM17 (B) expression in paclitaxel responders and non-responders in the TCGA dataset, showing higher expression in responders. (C–E) Analysis of FRMD8 (C), iRhom2 (D), and ADAM17 (E) expression in paclitaxel responders and non-responders in the GSE22513 dataset, supporting TCGA findings. (F) qPCR analysis of iRhom2 mRNA levels in 759 cells treated with paclitaxel, anastrozole, and doxorubicin, confirming increased iRhom2 with paclitaxel. (G) MTT assay assessing drug sensitivity to docetaxel, showing similar results to paclitaxel. (H) Gross images from the drug screening experiment, highlighting increased sensitivity of FRMD8 knockdown cells to paclitaxel. (I) In vivo imaging of GFP signals in primary tumors, showing reduced tumor growth with treatments. (J–P) Gross images, GFP signal measurements, and H&E staining of spleen (J), kidneys (K, L), liver (M, N), and brain (O, P) from treated mice, demonstrating effective inhibition of metastasis in multiple organs with combined treatment. Data are presented as mean ± SD (n = 5 mice per group). Statistical significance was assessed using a two-tailed Student’s t-test. ***P < 0.001.

